# Peripheral Neuropathy After Chronic Alcohol Exposure in Mice: Impact of sex, total intake and duration and alcohol metabolism

**DOI:** 10.1101/2025.09.02.673579

**Authors:** L.V. Moncayo, F. Adu-Gyamfi, A. Mohiuddin, M. Cruz, A. Dahman, Z. Akbar, S. Kidd, A. Siddiqi, A. Khan, A. Chiang, T. Singh, S. Herz, L. Mawaldi, G Valencia, P. Patel, A. Rauf, M.F. Miles, M.I Damaj

## Abstract

Alcohol induced peripheral neuropathy (AIPN) is a neurodegenerative disease caused by chronic alcohol intake and is associated with peripheral nerve damage and somatosensory symptoms, such as allodynia. Current treatments lack efficacy and do not target underlying pathology emphasizing the need for preclinical models of AIPN to elucidate mechanisms and novel targets. Thus, we performed a detailed characterization of a mouse model of AIPN and candidate mechanistic associations including the role of neuroinflammation and acetaldehyde. Our studies showed chronic alcohol induced mechanical and cold hypersensitivity and deficits in spontaneous behaviors in EtOH concentration-, time- and sex-dependent manners. Female mice drank more alcohol and developed more rapid and severe hypersensitivity but less robust deficits of spontaneous behaviors. The grimace test demonstrated chronic alcohol promoted spontaneous pain independent of sex. Duration of intake impacted alcohol-induced deficits in peripheral nerve electrophysiology amplitude and intra-epidermal nerve fiber density. We characterized an extensive time-course of chronic alcohol-induced neuroinflammation in the DRG and spinal cord and found significant time, sex and tissue effects. Polymorphisms of ALDH_2_ have been associated with alcohol-induced neuroinflammation, alcohol-related pain, and alcohol-induced peripheral neuropathy. We investigated the role of acetaldehyde, via inhibition of ALDH2, in the development of AIPN and showed that ALDH2 inhibition accelerated and exacerbated development of chronic alcohol-induced hypersensitivity in male and female mice. Overall, our studies in a well-controlled model of AIPN strongly point to neuroinflammation and inflammatory modulators such as acetaldehyde as important mechanistic targets for possible intervention in AIPN.

## A. INTRODUCTION

Alcohol use disorder (AUD) is a major public health burden in the United States (SAMHSA, 2023), and peripheral neuropathy is one of the common neurologic complications that is associated with AUD. Alcohol-induced peripheral neuropathy (AIPN) is a progressive and prevalent somatosensory disease by which individuals with a history of chronic alcohol use present disruptions in peripheral nerve integrity and debilitating neuropathic pain (Julian et al., 2019). A recent meta-analysis found that about 50% (95% CI, 35.9% to 53%) of patients with AUD have peripheral neuropathy (Julian et al., 2019) and more recent smaller studies report up to 60% of AUD patients were diagnosed with peripheral neuropathy (Papantoniou et al., 2024).

There is no reliable successful therapy for AIPN and the current pharmacotherapies (i.e., tramadol and gabapentin) poorly manage symptoms and have little impact on disease progression or recovery (Chopra & Tiwari, 2012). Consequently, there is a need for new agents for prevention and/or reversal of AIPN. The clinical investigation of AIPN presents several challenges which can confound mechanistic research, such as variability in the prevalence and extent of patient self-reported neuropathy endpoints as in somatosensory symptoms, difficulty in quantifying alcohol intake and the variability in nerve fiber disruption possibly due to concurrent disease or environmental factors. Thus, the use of pre-clinical models of AIPN provides opportunities to test specific hypothesis and risk factors, elucidate underlying pathophysiological mechanisms, and possibly identify novel pharmacological targets for AIPN. With pre-clinical alcohol-related pain research on the rise, it is important to properly characterize pre-clinical models of AIPN that are relevant to human conditions and experiences. However, many reported rodent AIPN studies do not take into consideration clinically relevant risk factors (i.e., duration and level of intake, prolonged periods of alcohol withdrawal and the use of female animals) on the development or recovery of AIPN. In addition, the majority of studies primarily rely on evoked behavioral endpoints, without evaluating the impact of chronic alcohol intake on spontaneous pain-related behaviors, peripheral nerve fiber integrity and other aspects of neuropathy. More recent studies that investigated the impact of such factors in chronic-alcohol related pain phenotypes found that some behavioral measures, such as somatosensory hypersensitivity, are sex-dependent and linked to total alcohol exposure (Alexander et al., 2023; Brandner et al., 2023).

Multiple mechanisms have been considered in AIPN, but the exact mediators remain unclear. The direct toxic role of alcohol and/or its metabolites mediating AIPN is a potential mechanism, Metabolism of alcohol by the enzyme alcohol dehydrogenase forms acetaldehyde, a well-established contributor to neuroinflammation, which is immediately further metabolized by aldehyde dehydrogenases (ALDH), such as ALDH2. Indeed, significant research suggests that acetaldehyde mediates some of the behavioral effects of alcohol (Arizzi-LaFrance et al., 2006; Correa et al., 2012; Peana & Acquas, 2013; Peana et al.,2008, 2009, 2016, 2017; Ledesma et al., 2013). While the role of acetaldehyde in the development of AIPN has yet to be thoroughly investigated, two small clinical studies elucidated human polymorphisms of Aldh2 that are associated with AIPN (Masaki et al., 2004; Yang et al., 2020). Additionally, a recent pre-clinical study demonstrated acetaldehyde generated by ADH in Schwann cells may initiate and sustain mechanical hypersensitivity in male mice via TRPV1-mediated mechanisms (De Logu et al., 2019). A role for pathways involving oxidative stress and inflammation has also been suggested. Existing literature also suggests that alcohol-induced neuroinflammation and oxidative stress contributes to the neurodegeneration associated with AIPN (Borgonetti et al., 2023; Chopra & Tiwari, 2012).

Therefore, this study aimed to use male and female C57BL/6J mice to 1) characterize the development and recovery of AIPN using comprehensive variety of dependent variables and endpoints, and 2) directly investigate the role of acetaldehyde in the development of AIPN. The impact of clinically relevant risk factors, such as alcohol concentration, duration of alcohol intake, prolonged alcohol withdrawal and sex on the development of AIPN were also evaluated.

## B. METHODS AND MATERIALS

### B.1. Animals

Adult male and female C57BL/6J mice (7-week-old at the start of the experiment) were acquired from Jackson Laboratory (Bar Harbor, ME, USA). Animals were housed in an animal care facility approved by the Association for the Assessment and Accreditation of Laboratory Animal Care (AALAC) 2 per cage at Virginia Commonwealth University (Richmond VA) on a regular 12-hour light/dark cycle. Mice were habituated to the animal care vivarium for one week prior to any change of diets or baseline measures taken. All experimental testing was performed during the light cycle. Up until animals were switched to the liquid diet, food and water were given ad libitum (7012 Teklad LM-485 Mouse/Rat Sterilizable Diet, Harlan Laboratories Inc., Indianapolis, IN). Animals were euthanized via decapitation for collection of tissues after testing was completed. All experiments were approved by the Institutional Animal Care and Use Committee at Virginia Commonwealth University. All mice were observed daily for general well-being and their weight was measured on a regular basis. The allocation to experimental groups of mice was randomized

### B.2 Drugs and administration

100% Biology Grade Ethanol was obtained from Sigma-Aldrich (St. Louis, MO). The selective aldehyde dehydrogenase-2 (ALDH-2) inhibitor, CVT10216, was acquired from Sigma-Aldrich (Cas #: 1005334-57-5) and was dissolved in 5% carboxy-methylcellulose (CMC) at a dose of 40 mg/kg via i.p. route, similar to doses found to increase acetaldehyde *in vivo* (Arolfo et al., 2021, Yu et al., 2024).

### B.3 Chronic alcohol drinking paradigm

Mice were fed alcohol Lieber-DeCarli liquid diets according to previously published literature with a slight modification (Bertola et al., 2013; De Logu et al., 2019). Mice were housed 2 per cage and fed 50 mL of control Lieber-DeCarli diet (F1259, BioServ) ad libitum for 7 days to acclimate them to eating out of the feeding tube and the liquid diet. Afterward, mice were separated, single housed and were given 30 mL of either a control Lieber-DeCarli liquid diet or an alcohol Lieber-DeCarli liquid diet. Based on previously published studies using the Lieber-Decarli diet for AIPN, alcohol liquid diets were either 5% (v/v) or 2.5% (v/v) ethanol (EtOH) depending on the experiment (De Logu et al., 2019). Control and alcohol liquid diets were calorically matched using maltose-dextrin. Mice were checked daily for health, alertness, body mass, and volume of liquid diet consumed. The volume of diet consumed was used to calculate the total amount of alcohol (mL) consumed daily. The volume of alcohol consumed (mL) was converted to the mass of EtOH consumed (g) by multiplying the volume consumed by the density of EtOH (0.79 g/mL). The EtOH consumed (g) was divided by the body mass (kg) daily to determine the amount of EtOH consumed relative to the body mass (g EtOH/kg body mass). The daily amount of EtOH consumed relative to body mass (g EtOH/kg body mass) for each given week of alcohol intake was averaged to calculate the average daily EtOH consumption for each week of alcohol intake. Mice remained on the liquid diet for one, two, four or six weeks depending on the study. For alcohol withdrawal studies, mice remained on their assigned liquid diet for four weeks and then switched back to normal pellet food and water bottles ad/libitum for ten weeks.

### B.4 Experimental design

One cohort of mice (n=10/sex/group) was used to evaluate the impact of chronic alcohol concentration (2.5% or 5% EtOH) on the development (one, two, three and four weeks of intake) and recovery (one, two, four, eight and ten weeks of withdrawal) of mechanical and cold hypersensitivity. The same animals were used to evaluate the impact of chronic alcohol concentration on nesting behavior after four weeks of alcohol intake. Testing was conducted from least to most invasive: (1) nesting, (2) mechanical hypersensitivity and (3) cold sensitivity. A second cohort of mice (n= 8/sex/group) was used to evaluate the impact of the duration of chronic alcohol (5% EtOH) intake (two, four and six weeks) on the development of spontaneous pain (grimace) and spontaneous behavioral changes: nesting, locomotor activity, wheel running, and burrowing. Again, testing was conducted from least to most invasive: (1) nesting, (2) locomotor activity, (3) grimace (4) burrowing and (5) wheel running. A third group of mice (n=10/sex/group) was used to evaluate the impact of duration of chronic alcohol (5% EtOH) intake (two or four weeks) on caudal nerve conduction electrophysiology. A fourth group of mice (n=8/sex/group) was used to evaluate the impact of the duration (two or four weeks) of chronic alcohol (5% EtOH) intake on intraepidermal nerve fiber (IENF) density. The same cohort of mice was used to evaluate the impact of duration (one, two, four or six weeks) of chronic alcohol (5% EtOH) intake on the differential RNA expression of pro-inflammatory cytokines in the DRG and spinal cord. A final cohort of animals (n= 5/sex/group) was used to evaluate the impact of ALDH2 inhibition on the development of AIPN.

### B.5 Mechanical hypersensitivity testing

Mechanical hypersensitivity was assessed using von Frey filaments. Mice were acclimated to the room for 30 minutes in their home cage before being placed in a Plexiglas cup where they were habituated on a mesh for 15 min prior to von Frey testing. Paw withdrawal (PW) thresholds were determined by applying a series of calibrated von Frey filaments using the up-down method as previously described (0.07-4.0 g of force) (Ghani et al., 2024). A PW was defined as immediate lifting, shaking or fluttering of the paw. The PW thresholds of both paws were averaged to provide a final PW threshold value. Mice were tested for baseline PW thresholds prior to alcohol exposure.

### B.6 Cold sensitivity testing

Cold sensitivity was assessed using an acetone test immediately following completion of mechanical hypersensitivity testing. 20 μL of acetone was applied to the right hind paw and the total PW time was measured for a 60 seconds after acetone application (Warncke et al., 2021). A PW was defined as a lift, flutter, or licking of the paw. Mice were tested for baseline PW times prior to alcohol exposure.

### B.7 Spontaneous Behavioral Battery

In addition to the evoked nociceptive hypersensitivity tests, we assessed spontaneous behavioral measures to reveal the presence of non-evoked ongoing pain. We used innate behaviors to assess the general well-being of mice by measuring the reduction in nesting behavior, burrowing behavior, hanging and rearing behaviors, and motor performance/motivation using the voluntary wheel running test as surrogate markers of pain-like behaviors. These tests employ evolutionarily conserved rodent behaviors suggested to represent behaviors that may represent the “activities of daily living” in humans (Wodarski et al., 2016). We also assessed general spontaneous locomotor activity.

#### Nesting

The nesting test evaluates the impact of spontaneous nociception on typical spontaneous activities in the daily mouse life. Nesting testing occurred in the same room that mice were housed, using the same: corncob bedding typically used in their home cages, nestlet material typically used in their home cages and their home cage lid. Nesting was assessed early in the morning during their peak nesting time (8-10 am) and was uninterrupted to ensure minimum disturbance during the testing period. Nesting testing occurred the day *prior* to typical cage change days. Nestlets were cut in half and 1-half nestlet piece was placed in each of the 4 corners of the testing cage. Mice were given 2 hours to nest and then pictures of the final nests built were taken. Pictures of the nesting cages were scored by a blinded observer on a scale of 1-5 to determine a “shred score” based on the amount the nestlets were brought together and how much the nestlets were shredded.

#### Hanging and Rearing

Mice were habituated to the behavioral apparatus for 30 min the day prior to testing. Mice were acclimated to the testing room 30 min prior to testing. Mice were placed in an empty mouse cage with a standard metal food hopper cover, and video footage was recorded using MED-PC IV software for 30 minutes. A blinded observer scored the videos to determine the total time (in minutes) the mice spent performing hanging or rearing behavior. Hanging behavior was defined as grabbing the metal hopper cover with a minimum of one paw. Rearing behavior was defined as placing weight on the hind paws while extending the torso towards the sky with the front paws. The percentage time spent hanging or rearing was calculated as [(total behavior time/30)*100].

#### Locomotor Activity

Mice were habituated to the testing room 30 minutes prior to locomotor activity assessment. Mice were individually placed in boxes (20 x 24 x 16.5 cm, Omnitech, Columbus, OH) and locomotor activity was recorded using MED-PC IV software for 120 minutes. Data expressed as the number of photobeam breaks during the testing session.

#### Burrowing

Similar to nesting, the burrowing test is another measure to assess spontaneous nociception-induced depression in spontaneous behaviors of mice. The burrowing protocol was adapted from previously described methods with some minor modifications (Warncke et al., 2021). Mice were habituated to the testing room 30 minutes prior to burrowing testing. A test cage was set up with a hollow tube, open on one end, and filled with 100 grams of corn cob bedding material. Mice were placed in a test cage for one hour with their own cage lid and allowed to burrow. The amount of corn cob bedding remaining at the end of the testing session was recorded. Data expressed as a percentage of the bedding removed.

#### Voluntary wheel running

The wheel running is another method to assess spontaneous nociception-induced depression in spontaneous behavior with an added motivational component. The wheel running test was performed as previously described (Contreras et al., 2021). Mice were placed in polycarbonate wheels (diameter 21.5 cm; width 5 cm) with a steel rod axle containing an electronic sensor which recorded the number of rotations in a session. The equipment for this assay was built locally (Virginia Commonwealth University, Richmond, VA, USA). Mice were habituated to the testing room 30 minutes prior to voluntary wheel running testing. The number of rotations in a 120-minute session was determined. Data expressed as total number of turns.

### B.8. Evaluation of Spontaneous Pain Grimace Score

Mice were evaluated for spontaneous pain using the grimace score according to previously published methods using a slightly modified protocol (McCoy et al., 2022). Mice were habituated to the testing room 30 min prior to grimace score evaluation. Mice were placed individually into lit behavioral chambers (Hypothesis to Hardware, LLC, Mouse Behavioral Chamber) directly in front of an HD camera and recorded for 15-min sessions. Video footage was uploaded to Painface Software (http://painface.net) (McCoy et al., 2024) for scoring using a medium frame speed. The grimace scores for the first 10 min (600 frames) were averaged and data expressed as an average grimace score.

### B.9. Caudal Nerve Conduction

Mice acclimated to the testing room at least one hour prior to nerve conduction assessment. Nerve conduction velocity and nerve action potential amplitudes of the caudal nerve were recorded with a PowerLab 4/35 electromyography system (AD Instruments, Inc., CO, USA) as previously described (Warncke et al., 2021). Mice were anesthetized individually in a chamber with 5% isoflurane carried with oxygen and maintained with 2.5% isoflurane by nose cone during the nerve conduction procedure. Cathode and anode needle recording electrodes were inserted 5 mm distal from the base of the tail, with the stimulating electrodes 5.0 com apart from the recording points (distance measured from cathode to cathode). A ground electrode was placed halfway between the stimulating and recording electrodes. The nerve was stimulated with single square-wave pulses of 0.1 ms duration at 7 mA, with a repeat rate of 1 Hz. This test measured the amplitude and latency of the evoked response and was used to determine the nerve conduction amplitude and velocity (0.05 m/latency). Mice were tested before alcohol exposure as a baseline measure and during alcohol intake. Data expressed as a change from baseline (baseline measure-timepoint measure).

### B.10 Immunohistochemistry and quantification of Intra-epidermal nerve fibers

Mouse hind paws were removed and the glabrous skin on the ventral surface of the hind paws were excised and submerged in freshly prepared periodate-lysine-paraformaldehyde (PLP) fixative at 4°C for 24 hours as previously described (Warncke et al., 2021). The tissues were then transferred to 30% (w/v) sucrose at 4° C within 48 hours. The tissues were then embedded in Optimal Cutting Temperature embedding medium for frozen tissues (ThermoFisher Scientific) and sectioned at 25 μM by cryostat. Sliced sections were dipped in cold acetone (−20°C) for 15 min, washed with phosphate-buffered saline (PBS) and incubated at room temperature for 45 minutes in blocking solution (5% normal goat serum and 0.3% Triton X-100 in PBS). Sections were then incubated with a diluted (1:200) primary rabbit anti-mouse polyclonal PGP9.5 antibody (ProteinTech, cat#14730-1-AP, IL, USA) overnight at 4°C. Following the overnight incubation, sections were washed 3 times with PBS and were incubated a second time in blocking solution prior to incubation for 2 hours at room temperature with diluted (1:300) goat anti-rabbit IgG (H+L) secondary antibody conjugated with Alexa Fluor® 594 (Life Technologies, cat# A11037, OR, USA). Sections were mounted with Vectashield (Vector Laboratories, Burlingame, CA, USA) and viewed using a Zeiss Axio Imager A1-Fluorescence microscope (Carl Zeiss, AG, Germany). The total number of IENFs for each paw section were counted under 40x magnification by a blinded quantifier and the mean IENF density was calculated from quantified sections. Data expressed as IENF density (fibers/mm).

### B.11 qRT-PCR

Snap frozen tissues (spinal cord overlaying the T13-L1 vertebrae and L4-6 lumbar dorsal root ganglia) were homogenized and purified RNA was extracted using the Invitrogen™ PureLink™ RNA Mini Kit (ThermoFisher scientific). Purified RNA concentrations for each sample were quantified using the Qubit 3.0 fluorimeter (ThermoFisher scientific). Extracted and purified RNA was diluted (15 ng/μL) and 15 μL was added to 5 μL of master mix [4 μL iScript 5x reaction mix, 1 μL reverse transcriptase; iScript™ cDNA Synthesis Kit (Biorad, Hercules)] to synthesize 20 μL of cDNA by reverse transcription. 4.4 μL of cDNA was then added to 39.6 μL of Taqman master mix (20 μL 2X SuperMix, 14 μL nuclease-free water, 2 μL random primer). mRNA expression levels of TNFα (Mm00443258_m1 Tnf, ThermoFisher scientific), IL-1β (Mm00434228_m1 Il1b, ThermoFisher scientific) and IL-6 (Mm00446190_m1 Il6, ThermoFisher scientific) were measured via quantitative real time polymerase chain reaction (qRT-PCR) assay. mRNA levels from all genes assessed were normalized using Beta-2-Microglobulin (B2M, Hs99999907_m1) as a reference gene. All qRT-PCR assays were performed using the TaqMan™ Gene Expression Master Mix (ThermoFisher scientific, Waltham, MA, USA), loaded on 96-well plates (Biorad, Hercules, CA, USA), and analyzed via a QuantStudio™ 3 Real-Time PCR System (ThermoFisher scientific, Waltham, MA, USA).

### B.12 Blood Acetaldehyde Quantification

Blood samples were collected retro-orbitally in male and female mice exposed 60 min after the 14^th^ administration of CVT10216 on the 14^th^ day of 2.5% EtOH intake for quantification of blood acetaldehyde levels. The whole blood acetaldehyde concentrations were determined using a modified method routinely employed for the analysis of clinical, forensic and other samples for ethanol, and volatiles (1). In brief, an 8-point calibration curve (0.8-160 mg/L acetaldehyde (Thermo Scientific Chemicals, Waltham, Mass)), controls consisting of Ethanol 80 control and Volatile controls (UTACK® Laboratories, Inc., Valencia, CA) along with blank controls with and without internal standard were prepared. Aliquots of one hundred microliters of calibrator, controls and samples were added to 900 µL of 234 mg/L n-propanol working internal standard (ISTD) in a 20 mL headspace vial. The samples analysis was then performed using Tekmar 7000 headspace autosampler with a Hydroguard MXT® 2 m × 0.53 mm ID (Restek Corp., Bellefonte, PA) sample loop and a Restek guard column attached to Varian 3900 gas chromatograph with an FID detector (Varian Associates, Inc., Walnut Creek, CA, USA) equipped with a Rtx®–BAC2 30 m × 0.53 mm ID × 2.0 μm column (Restek Corp., Bellefonte, PA). The oven and detector temperatures were 40 and 235°C, respectively with nitrogen used as the carrier gas. Calibration curves were constructed for acetaldehyde using linear regression of peak area ratios of the analyte to the ISTD.

### B.13 Statistical Analysis

Data obtained were analyzed using GraphPad software (GraphPad Software 9.2, Inc., La Jolla, CA) or RStudio and are expressed as the mean ± SEM. All data passed normality (Shapiro-Wilk test) and equal variance (Brown-Forsythe test). To determine the impact of sex as a biological variable, data was first analyzed and reported using a 1-, 2-, or 3-Way ANOVA (See Supplemental Tables). If sex was determined not to be a significant factor, male and female data were pooled, and sex was collapsed as a factor; results were then analyzed and reported as mixed sex. If sex was revealed to be a significant factor, male and female data were further analyzed separately. Follow up analysis (male, female or mixed sex) was conducted using the Un-Paired Student t-test, 1-Way, 2-Way, and 3-Way ANOVA tests and were followed by Tukey’s post hoc analysis when appropriate. Pearson correlations were used to determine the relationship and significance between endpoints in studies and total EtOH consumption, and this data is reported as the Pearson coefficient (R), followed by the p-value.

## C. RESULTS

### Alcohol concentration and sex influence EtOH intake

Mice were exposed to two concentrations of alcohol (2.5 and 5% EtOH) for four weeks (**Figure 1A**). Alcohol concentration, time and sex impacted the average daily EtOH intake in C57BL/6J mice. Throughout the study, female mice subjected to a higher concentration, consumed more EtOH compared to those on a lower concentration [F_(1, 14)_ = 103.5, p<0.0001, **Figure 1B**]. In male mice, a time-dependent effect of alcohol concentration on daily EtOH intake was observed [F_(3, 40)_ = 12.18, p<0.0001]. Male mice exposed to 5% EtOH diets consumed higher levels of EtOH, compared to mice on 2.5%, that gradually increased over time [F_(3, 40)_ = 12.18, p<0.0001, **Figure 1C**]. Tukey’s post-hoc analysis revealed male mice on 5% EtOH diets consumed more EtOH at 1 (P<0.001), 2 (P<0.0001), 3 (P<0.0001) and 4 (P<0.0001) weeks. Analysis of the total EtOH consumed throughout the four weeks of alcohol intake (**Figure 1D**) revealed a significant sex-dependent effect of alcohol concentration [F_(1, 28)_ = 8.941, p=0.0058]. In both males (P<0.0001) and females (P<0.0001), mice on 5% EtOH diets consumed more EtOH compared to mice on 2.5% EtOH diets. For mice given 5% EtOH diets, female mice consumed more EtOH than male mice (P<0001). Analysis of the average weekly body mass over the four weeks of alcohol intake revealed sex as a significant factor (**Supplementary Table 1**), as female body masses were lower than male mice [F_(1, 166)_ = 110.6, p<0.0001]. In general, female mice on alcohol diets showed lower body mass than control diet mice throughout the experiment [F_(2,21)_ = 7.266, p=0.004, **Figure 1E**]. Male mice exposed to alcohol displayed attenuated weight gain over time [F_(2, 21)_ = 11.29, p=0.0005; F_(6, 61)_ = 2.873, p=0.0157; respectively, **Figure 1F**], and by four weeks of alcohol intake, the effect in mice given 5% EtOH was significantly more severe than those given 2.5% (P<0.05).

**Figure 1.**
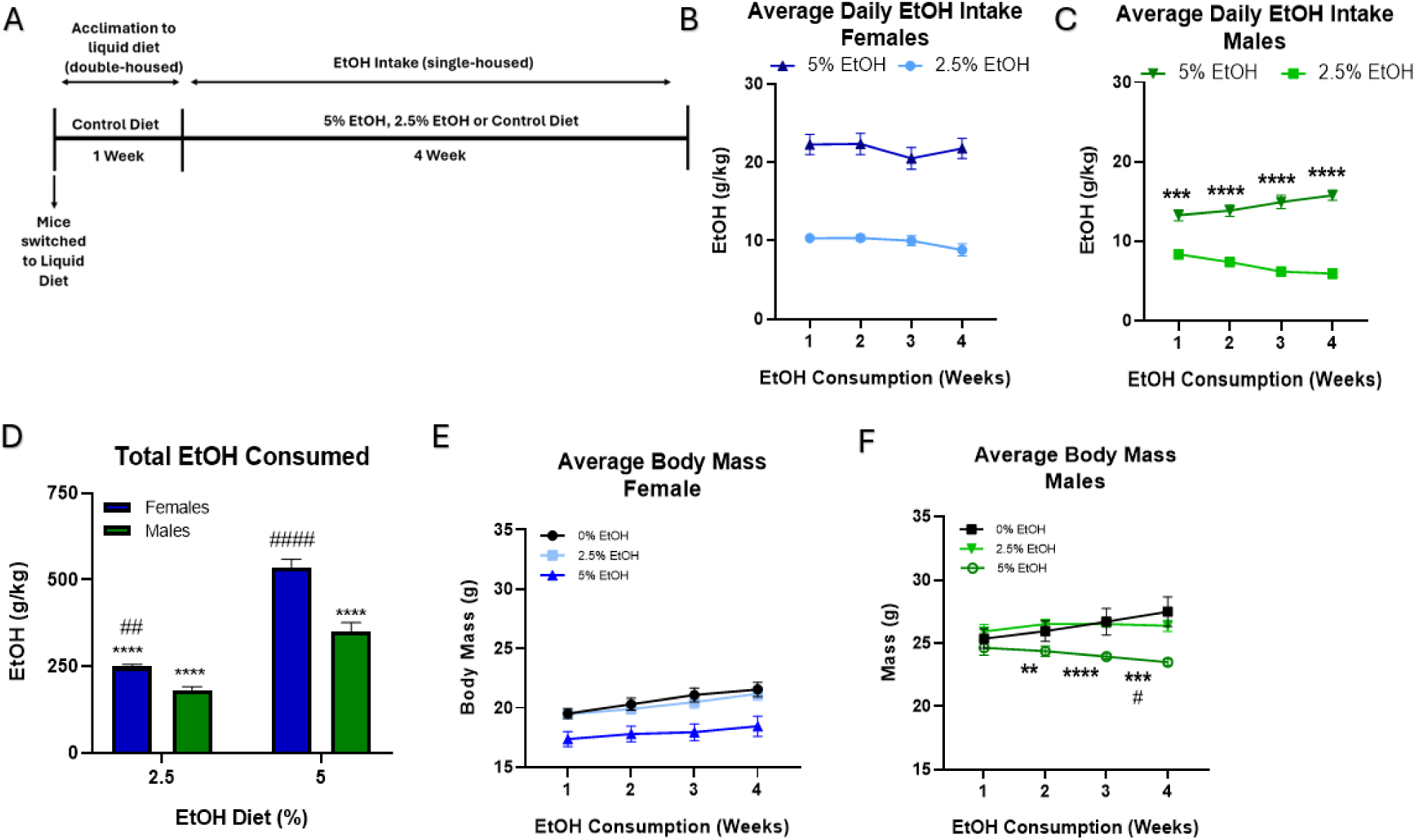
│ The effect of EtOH concentration on EtOH consumption and body mass over four weeks in C57BL/6J mice (8/sex/group) Values expressed as mean ± SEM. **(A)** Timeline of alcohol liquid diet paradigm. **(B)** Average daily EtOH intake in female mice. Results were compared using two-way ANOVA (EtOH % x Time as RM). **(C)** Average daily EtOH intake in male mice. Results were compared using two-way ANOVA (EtOH % x Time as RM) followed by Tukey’s post-hoc test, ***<0.001 versus 2.5% EtOH, ****p<0.0001 versus 2.5% EtOH. **(D)** Total EtOH consumed in male and female mice. Results were compared using two-way ANOVA (EtOH % x Sex) followed by Tukey’s post-hoc test, ****p<0.0001 versus 5% female, ## p<0.01 versus 5% males, ####p<0.0001 versus 5% males. **(E)** Average daily body mass of female mice. Results compared via two-way ANOVA (EtOH % x time as RM). **(F)** Average daily body mass of male mice. Results compared via two-way ANOVA (EtOH % x time as RM) followed by Tukey’s post-hoc test, **p<0.01 versus 0% EtOH control diet, ***p<0.001 versus

### Chronic alcohol induces long-term, concentration- and sex-dependent mechanical hypersensitivity

We evaluated the impact of the two concentrations of alcohol (2.5 and 5% EtOH) on the time-course of stimulus-evoked nociception related behaviors during four weeks of alcohol intake and ten weeks after alcohol cessation (Figure 2A). No significant difference in paw withdrawal threshold was observed between groups prior to alcohol exposure. In both male and female mice, A Two-Way ANOVA (sex x EtOH %) analysis of the area under the curve revealed a main effect of EtOH concentration on the development [F_(2, 231)_ = 18.48, p<0.0001, **Supplemental Figure 1A**] and recovery [F_(2, 270)_ = 63.22, p<0.0001, **Supplemental Figure 1B**] of mechanical hypersensitivity. Additionally, chronic alcohol intake induced long-term mechanical hypersensitivity in a concentration-, time- and sex-dependent manner [F_(3,82)_ = 4.262, p=0.0075, Supplemental Table 2].

**Figure 2.**
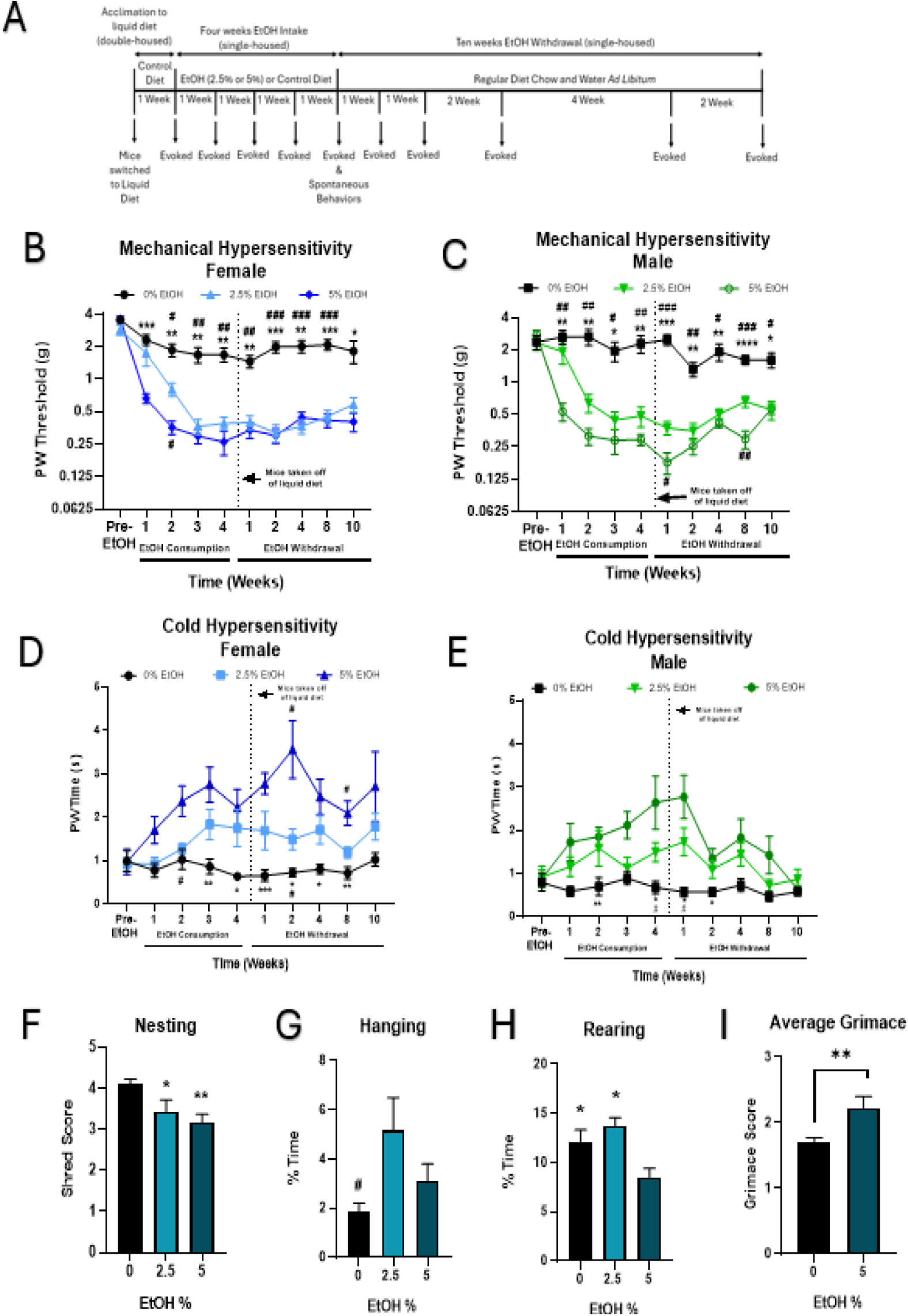
│ The effect of EtOH concentration on the development of mechanical and cold hypersensitivity and nesting behavior over four weeks in C57BL/6J mice. Values expressed as mean ± SEM. **(A)** Timeline of alcohol liquid diet paradigm and behavioral testing. **(B)** EtOH concentration-dependent development of mechanical hypersensitivity in female mice. (n=8/group). Results were compared using two-way ANOVA (EtOH % x Time as RM) followed by Tukey post-hoc test, *p<0.05 versus 5% EtOH, **p<0.01 versus 5% EtOH, ***p<0.001 versus 5% EtOH, #p<0.05 versus 2.5% EtOH, ##p<0.01 versus 2.5% EtOH, ###p<0.001 versus 2.5% EtOH. **(C)** EtOH concentration-dependent development of mechanical hypersensitivity in male mice. (n=8/group). Results were compared using two-way ANOVA (EtOH % x Time as RM) followed by Tukey post-hoc test, *p<0.05 versus 5% EtOH, **p<0.01 versus 5% EtOH, ***p<0.001 versus 5% EtOH, ****p<0.0001 versus 5% EtOH, #p<0.05 versus 2.5% EtOH, ##p<0.01 versus 2.5% EtOH, ###p<0.001 versus 2.5% EtOH. **(D)** EtOH concentration-dependent development of cold hypersensitivity in female mice. (n=8/group). Results were compared using two-way ANOVA (EtOH % x Time as RM) followed by Tukey post-hoc test, *p<0.05 versus 5% EtOH, **p<0.01 versus 5% EtOH, ***p<0.001 versus 5% EtOH, #p<0.05 versus 2.5% EtOH. **(E)** EtOH concentration-dependent development of mechanical hypersensitivity in male mice. (n=8/group). Results were compared using two-way ANOVA (EtOH % x Time as RM) followed by Tukey post-hoc test, *p<0.05 versus 5% EtOH, **p<0.01 versus 5% EtOH, ***p<0.001 versus 5% EtOH, #p<0.05 versus 2.5% EtOH. **(F)** Alcohol induced nesting deficits in male and female mice (n=8-12/sex/group). Results compared via one-way ANOVA followed by Tukey’s post-hoc test, *p<0.05 versus 0% EtOH control diet, **p<0.01 versus 0% EtOH control diet. **(G)** Impact of chronic EtOH concentration on spontaneous hanging behavior in male and female mice (n=6-10/sex/group). Results compared via one-way ANOVA followed by Tukey’s post-hoc test, #p<0.05 versus 2.5%. **(H)** Impact of chronic EtOH concentration on spontaneous hanging behavior in male and female mice (n=6-10/sex/group). Results compared via one-way ANOVA followed by Tukey’s post-hoc test, *p<0.05 versus 5%. **(I)** Chronic alcohol-induced increase in grimace score in male and female mice (n=7-14/sex/group). Results compared via Un-Paired Student T-Test **p<0.01

In female mice, both alcohol concentrations caused reductions in paw withdrawal threshold over the course of alcohol intake that persisted throughout ten weeks after alcohol withdrawal [F_(18, 183)_ = 2.726, p=0.004, Figure 2B]. Tukey’s post hoc analysis revealed female mice exposed to 2.5% EtOH had lower PW thresholds compared to control diet mice at 2 (p<0.05), 3 (p<0.01) and 4 (p<0.01) weeks of alcohol intake and 1 (p<0.01), 2 (p<0.001), 4 (p<0.001) and 8 (p<0.001) weeks after alcohol withdrawal. Exposure to 5% EtOH induced quicker, more robust and persistent mechanical hypersensitivity compared to 2.5% EtOH. Female mice on 5% EtOH had lower PW thresholds than control mice at 1 (p<0.001), 2 (p<0.01), 3 (p<0.01) and 4 (p<0.01) weeks of EtOH intake. This reduction was still observed at 1 (p<0.01), 2 (p<0.001), 4 (p<0.01), 8 (p<0.001), and 10 (p<0.05) weeks after alcohol withdrawal. Additionally, a significant difference in PW threshold between 5% and 2.5% was observed at week two of alcohol intake (p<0.05) in female mice. Although Pearson correlation analysis between the total EtOH female mice consumed and the percent of their baseline PW threshold after 4 weeks of alcohol intake revealed a significant negative correlation (R=0.2715, p=0.0464, **Supplemental Figure 1E**), the negative correlation between the total EtOH consumed and the PW threshold after ten weeks of alcohol withdrawal was not significant (R=0.1878, p=0.1066, **Supplemental Figure 1G**).

A significant three-way interaction confirmed that EtOH concentration induced long-term mechanical hypersensitivity in male mice [F_(18, 182)_ = 4.357, p<0.0001, **Figure 2C**). Male mice exposed to chronic 2.5% EtOH had lower PW thresholds compared to control diet mice at 1 (p<0.01), 2 (p<0.01), 3 (p<0.05) and 4 (p<0.01) weeks of alcohol intake. This reduction in paw withdrawal threshold persisted despite 1 (p<0.001), 2 (p<0.01), 4 (p<0.05), 8 (p<0.001) and 10 (p<0.05) weeks of alcohol withdrawal. Similarly, 5% EtOH reduced PW thresholds at 1 (p<0.01), 2 (p<0.01), 3 (p<0.05) and 4 (p<0.01) weeks of alcohol intake in male mice. Tukey’s post-Hoc analysis further revealed 5% EtOH-induced reductions in paw withdrawal threshold were still significant at 1 (p<0.001), 2 (p<0.01), 4 (p<0.01), 8 (p<0.0001), and 10 (p<0.05) weeks of alcohol withdrawal. Interestingly, 5% EtOH caused a more persistent degree of mechanical hypersensitivity compared to 2.5% at 1 (p<0.05) and 8 weeks (p<0.01) of alcohol withdrawal. Similar to female mice, Pearson correlation analysis revealed a significant negative correlation (R=0.2715, p=0.0469, **Supplemental Figure 1F**) between the total EtOH male mice consumed and the percent of their baseline PW threshold after 4 weeks of alcohol intake. The correlation between the total EtOH consumed and the PW threshold after ten weeks of alcohol withdrawal was not significant and not as robust as that of female mice (R=0.01377, p=0.6770, **Supplemental Figure 1H**).

### Chronic alcohol induces cold hypersensitivity in a time-, concentration- and sex-dependent manner

In both male and female mice, 2-Way ANOVA (sex x EtOH %) of the area under the curve revealed a main effect of EtOH concentration on the development [F_(2, 226)_ = 9.266, p<0.0001, **Supplemental Figure 1C**] and recovery [F_(2, 208)_ = 28.29, p<0.0001, **Supplemental Figure 1D**] of cold hypersensitivity. Additionally, three-way ANOVA revealed a significant 2-way interaction (sex x time) impacted the development and recovery of chronic alcohol -induced cold hypersensitivity [F_(9, 288)_ = 2.978, p=0.0021, **Supplemental Table 2**].

In female mice, both alcohol concentrations caused the development of persistent cold hypersensitivity over time [F_(18, 177)_ = 1.918, p=0.0170; **Figure 2D**]. Tukey’s post hoc analysis determined PW times of female mice exposed to chronic 2.5% EtOH were significantly higher than control diet mice at 2 weeks of alcohol intake (p<0.05). However, the cold hypersensitivity partially recovered, and female mice given 2.5% EtOH were only significantly different than control diet mice at 2 (p<0.05) weeks of alcohol withdrawal. Female mice given chronic 5% EtOH had higher PW times at 3 (p<0.01) and 4 (p<0.05) weeks of alcohol intake. Interestingly, PW thresholds of female mice on 5% EtOH were still significantly higher at 1 (p<0.001), 2 (p<0.05), 4 (p<0.05) and 8 (p<0.01) weeks of alcohol withdrawal. Additionally, the higher concentration of alcohol caused significantly higher PW times than the lower concentration in female mice at 2 (p<0.05) and 8 (p<0.05) weeks of alcohol withdrawal. In male mice, chronic alcohol induced transient cold hypersensitivity in a concentration-dependent manner [F_(18, 175)_ = 1.872; p=0.0208; **Figure 2E**]. Tukey’s post hoc analysis showed chronic 2.5% EtOH caused higher PW times after 4 (p<0.05) weeks of alcohol intake. However, this increase in PW time was short-lasting, and mice exposed to 2.5% EtOH fully recovered. A more rapid onset of cold sensitivity was observed in male mice exposed to 5% EtOH. Paw withdrawal times were significantly increased at 2 (p<0.01) and 4 (p<0.05) weeks of alcohol intake in male mice given chronic 5% EtOH. Similarly, male mice given chronic 5% EtOH diets also fully recovered from cold hypersensitivity during an extended period of alcohol withdrawal. PW times of male mice on 5% EtOH were only significantly different from control diet mice at 1 (p<0.001) week of alcohol withdrawal.

### Chronic alcohol intake impacts spontaneous behaviors in a concentration-dependent manner

Four weeks of chronic alcohol reduced nesting behavior in a concentration-dependent but sex-independent manner [**Supplemental Table 2**]. One-way ANOVA revealed alcohol concentration as a significant factor in chronic alcohol-induced nesting deficits in female and male mice [F_(2, 59)_ = 4.933, p=0.0014, **Figure 2F**]. Further Tukey’s post hoc analysis determined female and male mice given control diet had higher nesting scores compared to mice exposed to chronic 2.5% (p<0.05) and 5% (p<0.01) EtOH. Additionally, four weeks of chronic alcohol impacted hanging behavior in a concentration-dependent but sex-independent manner [**Supplemental Table 2**]. One-way ANOVA revealed alcohol concentration as a significant factor in the impact of chronic alcohol on hanging behavior in female and male mice [F_(2, 41)_ = 2.143, p=0.0170, **Figure 2G**]. Further Tukey’s post hoc analysis determined female and male mice given 2.5% EtOH diets spent more time hanging compared to mice exposed to control diet (p<0.05). Alternatively, four weeks of chronic alcohol reduced rearing behavior in a concentration-dependent but sex-independent manner [**Supplemental Table 2**]. One-way ANOVA revealed alcohol concentration as a significant factor in chronic alcohol-induced nesting deficits in female and male mice [F_(2, 43)_ = 4.933, p=0.0484, **Figure 2H**]. Further Tukey’s post hoc analysis determined female and male mice given control diet had higher nesting scores compared to mice exposed to chronic 2.5% (p<0.05) and 5% (p<0.01) EtOH.

### Chronic alcohol induces spontaneous pain

The impact of chronic alcohol on the facial grimace pattern of mice was evaluated after six weeks of alcohol exposure. 2-Way ANOVA analysis determined chronic alcohol impacted grimace score in a concentration-dependent manner, independent of sex [**Supplemental Table 2**]. In both male and female mice, chronic 5% EtOH caused an increase in the average grimace score [t_(33)_ = 3.165, p=0.0033, **Figure 2I**].

### Chronic alcohol-induced spontaneous behavioral deficits are dependent upon time and sex

The impact of chronic alcohol intake on spontaneous behaviors (nesting, locomotor activity, burrowing, and wheel running) was evaluated in C57BL/6J mice following 2, 4 and 6 weeks of alcohol intake (**Figure 3A**). The higher concentration of 5% EtOH was selected based on the effects observed in the earlier nesting study.

**Figure 3.**
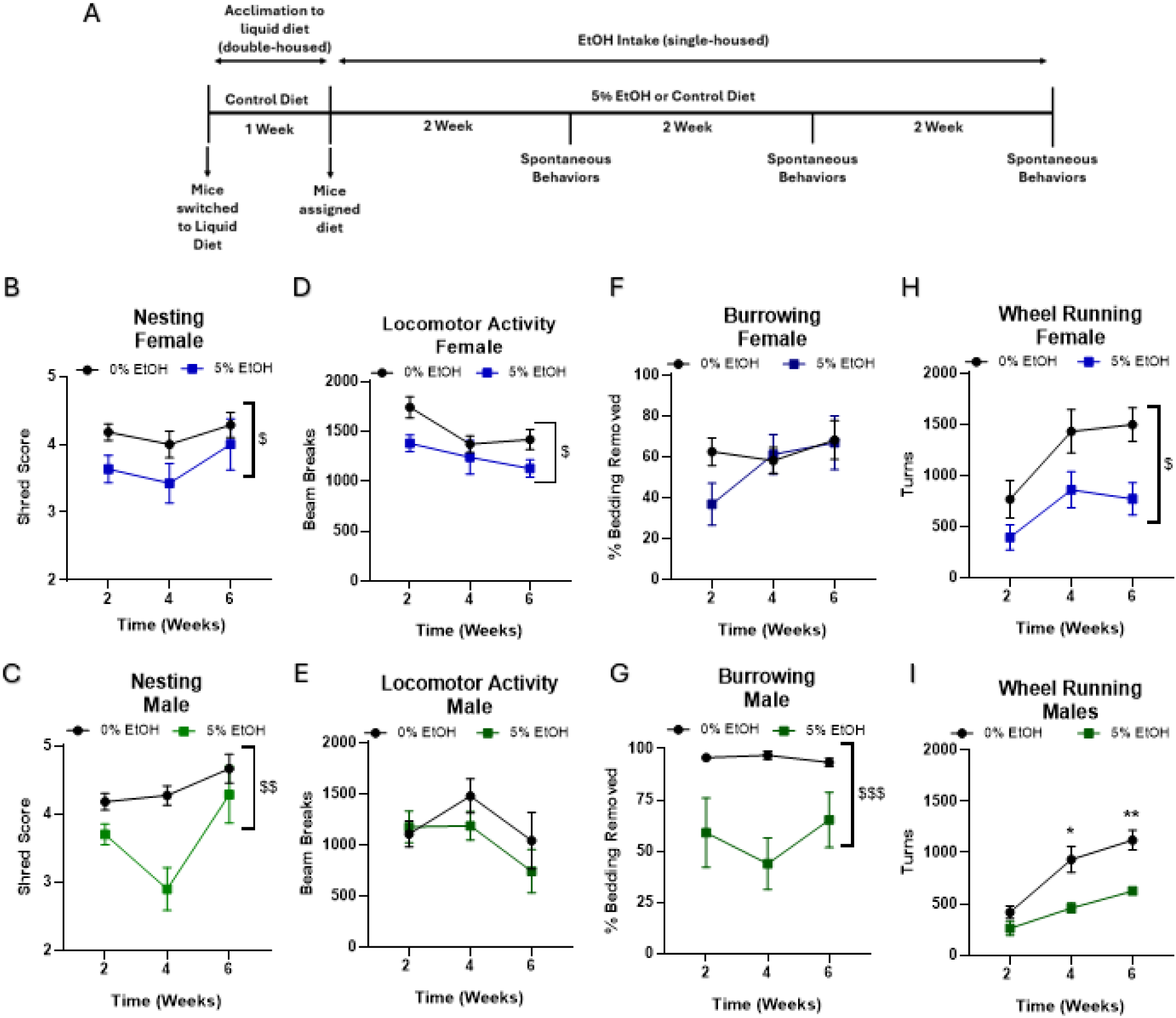
│The effect of duration of alcohol intake on spontaneous pain and spontaneous behaviors over six weeks in C57BL/6J mice. Values expressed as mean ± SEM. (n=8-15/sex/group). **(A)** Timeline of alcohol liquid diet paradigm and behavioral testing. **(B)** Chronic alcohol-induced nesting deficits in female mice. Results were compared using two-way ANOVA (EtOH % x Time as RM), $p<0.05 Main Effect of EtOH% **(C)** Chronic alcohol-induced nesting deficits in male mice. Results were compared using two-way ANOVA (EtOH % x Time as RM), $$p<0.01 Main Effect of EtOH% **(D)** Chronic alcohol-induced reductions of locomotor activity in female mice. Results were compared using two-way ANOVA (EtOH % x Time as RM), $p<0.05 Main Effect of EtOH% **(E)** Effects of chronic alcohol intake on locomotor activity in male mice. Results were compared using two-way ANOVA (EtOH % x Time as RM). **(F)** Effects of chronic alcohol intake on burrowing behavior in female mice. Results were compared using two-way ANOVA (EtOH % x Time as RM). **(G)** Chronic alcohol-induced deficits in burrowing behavior in male mice. Results were compared using two-way ANOVA (EtOH % x Time as RM), $$$p<0.001 Main Effect of EtOH %. **(H)** Chronic alcohol-induced deficits in wheel running behavior in female mice. Results were compared using two-way ANOVA (EtOH % x Time as RM), $p<0.05 Main Effect of EtOH %. **(I)** Chronic alcohol-induced deficits in wheel running behavior in female mice. Results were compared using two-way ANOVA (EtOH % x Time as RM) followed by Tukey Post hoc test, *p<0.05 versus 0% EtOH, **p<0.01 versus 0% EtOH

### Chronic alcohol impacts spontaneous behaviors in a time- and sex-dependent manner

Mice were evaluated in a battery of spontaneous behaviors following two, four and six weeks of alcohol intake. Unlike previously observed, 3-Way ANOVA revealed a significant main effect of sex on nesting behavior when evaluated at several time points [F_(1, 48)_ = 15.81, p=0.0002, **Supplemental Table 3**]. Although 2-Way ANOVA determined there was not a significant interaction of time and EtOH Diet % [F_(2, 31)_ = 0.2044, p=0.8162) or main effect of time [F_(1.877, 29.09)_ = 1.807, p=0.1817) on nesting behavior in female mice, a main effect of EtOH Diet % was observed [F_(1, 28)_ = 4.463, p=0.0437]. Overall, female mice exposed to 5% EtOH diets had lower shred scores compared to those exposed control diets (**Figure 3B**). Similarly in male mice, there was not a significant interaction of time and EtOH Diet % [F_(2, 29)_ = 3.173, p=0.0567]. However, there was a main effect of time [F_(1.303, 18.89)_ = 6.979, p=0.0111] and as expected, a main effect of EtOH Diet % [F_(1, 20)_ = 13.40, p=0.0016]. In general, male mice exposed to 5% EtOH diets had lower shred scores compared to those on control diets (**Figure 3C**).

Analysis via 3-Way ANOVA revealed an interaction of time and sex on locomotor activity [F_(2, 40)_ = 8.124, p=0.0011, **Supplemental Table 3**). In female mice, there was not a time-dependent effect of alcohol on locomotor activity [F_(2, 24)_ = 1.342, p=0.2802]. As expected, locomotor activity increased over time as female mice became acclimated to experimentation [F_(1.939, 23.27)_ = 8.510; p=0.0018). Additionally, female mice exposed to chronic alcohol had lower locomotor activity overall [F_(1, 13)_ = 5.160, p=0.0408, **Figure 3D**]. Neither this effect of chronic alcohol [F_(1, 14)_ = 0.7001, p=0.4168], nor an interaction of alcohol and time [F_(2, 16)_ = 2.063, p=0.1596], was found to impact locomotor activity in male mice. Locomotor activity did not change in male mice throughout experimentation [F_(1.580, 12.64)_ = 3.358, p=0.0761, **Figure 3E**].

Burrowing behavior was found to be impacted by chronic alcohol in only male mice [F_(1, 42)_ = 5.554, p=0.0232, **Supplemental Table 3**]. In female mice chronic alcohol exposure did not impact burrowing behavior [F_(2, 28)_ = 1.211, p=0.3131, **Figure 3F**]. Burrowing behavior was not impacted by time [F_(1.661, 23.25)_ = 0.5260, p=0.5650] or alcohol [F_(1, 21)_ = 0.8973, p=0.3543] in female mice. Similarly in male mice, 2-Way ANOVA did not reveal a significant interaction of time by alcohol [F_(2, 47)_ = 0.6861, p=0.5085] or main effect of time [F_(1.080, 25.37)_ = 0.3703, p=0.5644] on burrowing behavior. However, chronic alcohol caused an overall decrease in burrowing behavior in male mice [F_(1, 47)_ = 17.97, p=0.001, **Figure 3G**]. Overall, sex significantly impacted wheel running behavior in mice [F_(1, 28)_ = 7.918, p=0.0089, **Supplemental Table 3**]. In female mice chronic alcohol did not impact the wheel running behavior in a time dependent manner [F_(2, 22)_ = 1.698, p=0.2061]. Wheel running behavior increased overall over time as female mice became acclimated to the apparatus [F_(1.946, 21.41)_ = 21.29, p<0.0001]. More importantly, chronic alcohol exposure caused an overall lower number of turns in female mice [F_(1, 14)_ = 7.695, p=0.0149, **Figure 3H**]. Alternatively, 2-Way ANOVA revealed a significant time-dependent impact of chronic alcohol on wheel running behavior in male mice [F_(2, 23)_ = 3.655; p=0.0418]. Chronic 5% EtOH significantly prevented an increase in wheel running behavior over time in male mice (**Figure 3I**). Post hoc analysis determined male mice had a significantly lower number of turns at four (p<0.05) and six (p<0.01) weeks of alcohol intake in male mice.

### Chronic alcohol impacts the amplitude of caudal nerve electrophysiology

Caudal nerve conduction of male and female C57BL/6J mice were measured to determine the functional impact of chronic 5% EtOH intake on peripheral nerves following 2 and 4 weeks of alcohol intake (**Figure 4A**). Prior to alcohol exposure, there were no statistically significant differences between groups in amplitude or velocity. Analysis via 3-Way ANOVA determined sex did not influence amplitude or velocity, or the impact of alcohol on amplitude or velocity (**Supplemental Table 4**). In male and female mice, 2-Way ANOVA analysis determined there was not an interaction of alcohol and time on the change of caudal nerve conduction amplitude [F_(1,40)_ = 0.05824, p=0.8105]. Chronic alcohol impacted the change of caudal nerve conduction amplitude in male and female mice. Overall, mice exposed to chronic 5% EtOH saw a larger reduction in caudal nerve conduction amplitude [F_(1, 43)_ = 4.830, p=0.0334, **Figure 4B**].

**Figure 4.**
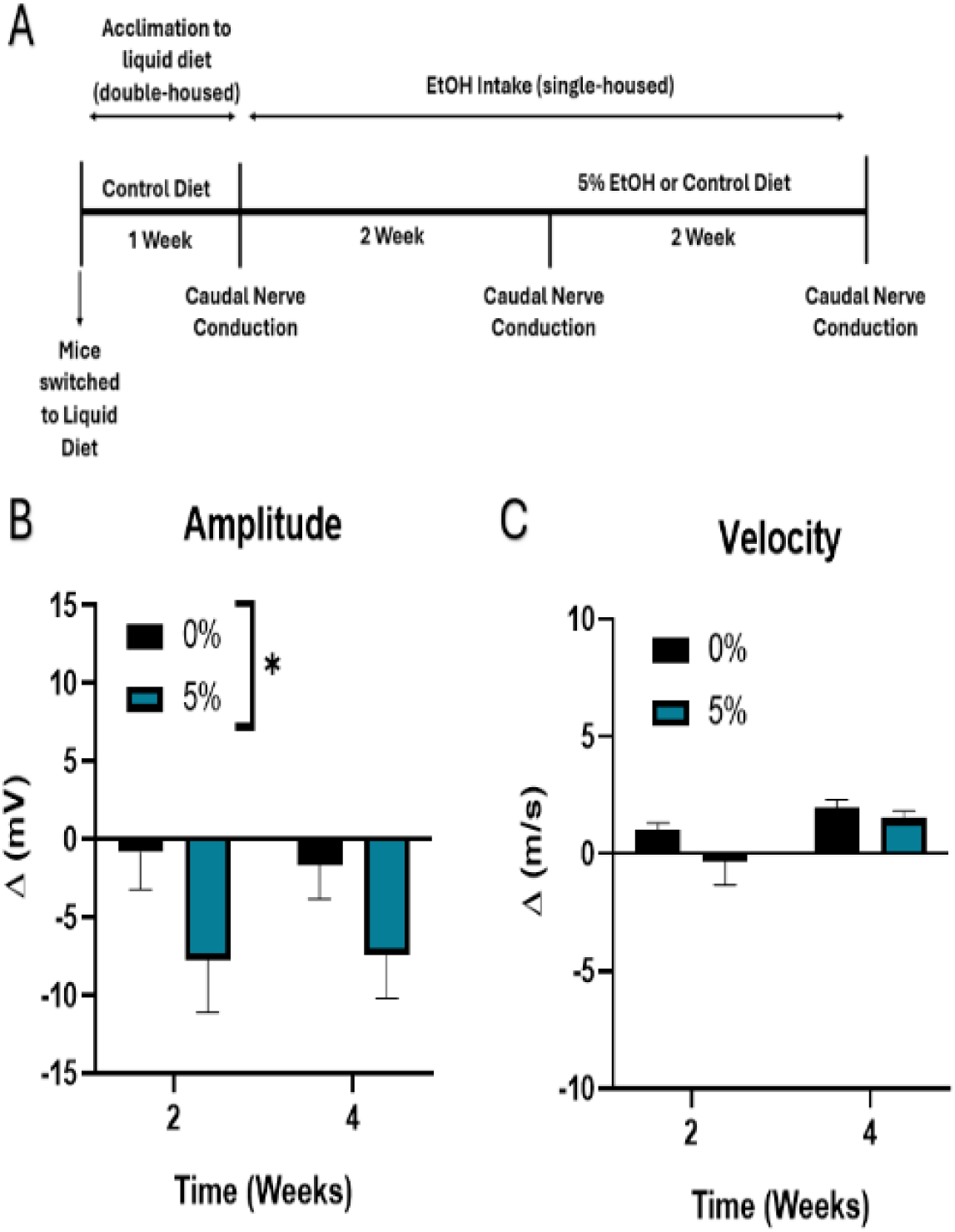
│ The effect of duration of alcohol intake on the change of caudal nerve conduction amplitude and velocity from baseline over four weeks in male and female C57BL/6J mice. Values expressed as mean ± SEM. (n=11-13/sex/group). **(A)** Timeline of alcohol liquid diet paradigm and caudal nerve conduction testing. **(B) C**hronic alcohol-induced deficits in caudal nerve conduction amplitude from baseline. Results were compared using two-way ANOVA (EtOH % x Time as RM), *p<0.05 Main Effect of EtOH %. **(C)** Effects of EtOH on the change of caudal nerve conduction velocity from baseline. Results were compared using two-way ANOVA (EtOH % x Time as RM).

Alternatively, 2-Way ANOVA determined only a main effect of time on the change in caudal nerve conduction velocity in male and female mice [F_(1, 43)_ = 6.015, p=0.0183, **Figure 4C**].

### Chronic alcohol reduced IENF density in male and female mice

**Supplemental Figure 2.**
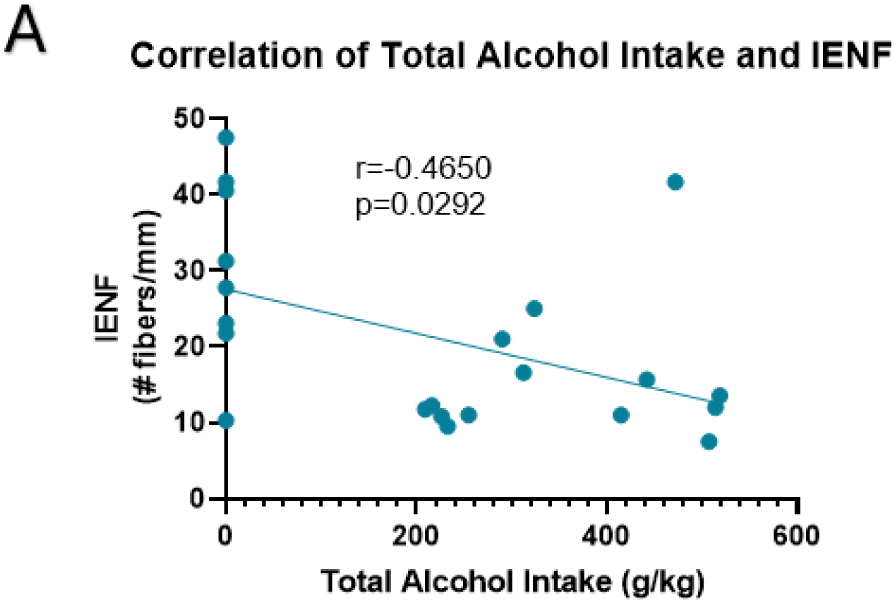
│Correlation of EtOH intake on IENF density in C57BL/6J mice (3-8/sex/group) (A) Pearson correlation between lifetime EtOH intake on IENF density in male and female mice exposed to chronic alcohol

The density of IENF was evaluated in groups of mice exposed to control diet, two weeks of 5% EtOH diet and four weeks of 5% EtOH diet (**Figure 5**). 2-Way ANOVA determined sex was not a significant factor on IENF density (**Supplemental Table 5**). In both male and female mice, the duration of alcohol intake impacted the IENF density [F_(2, 36)_ = 4.918, p<0.0001]. Compared to mice exposed to control diet, mice exposed to two (p<0.0001) or four weeks of 5% EtOH diets displayed lower IENF counts (p<0.0001). Pearson correlation analysis determined the total alcohol intake negatively correlated with the IENF density in male and female mice [**Supplemental Figure 2A**].

**Figure 5.**
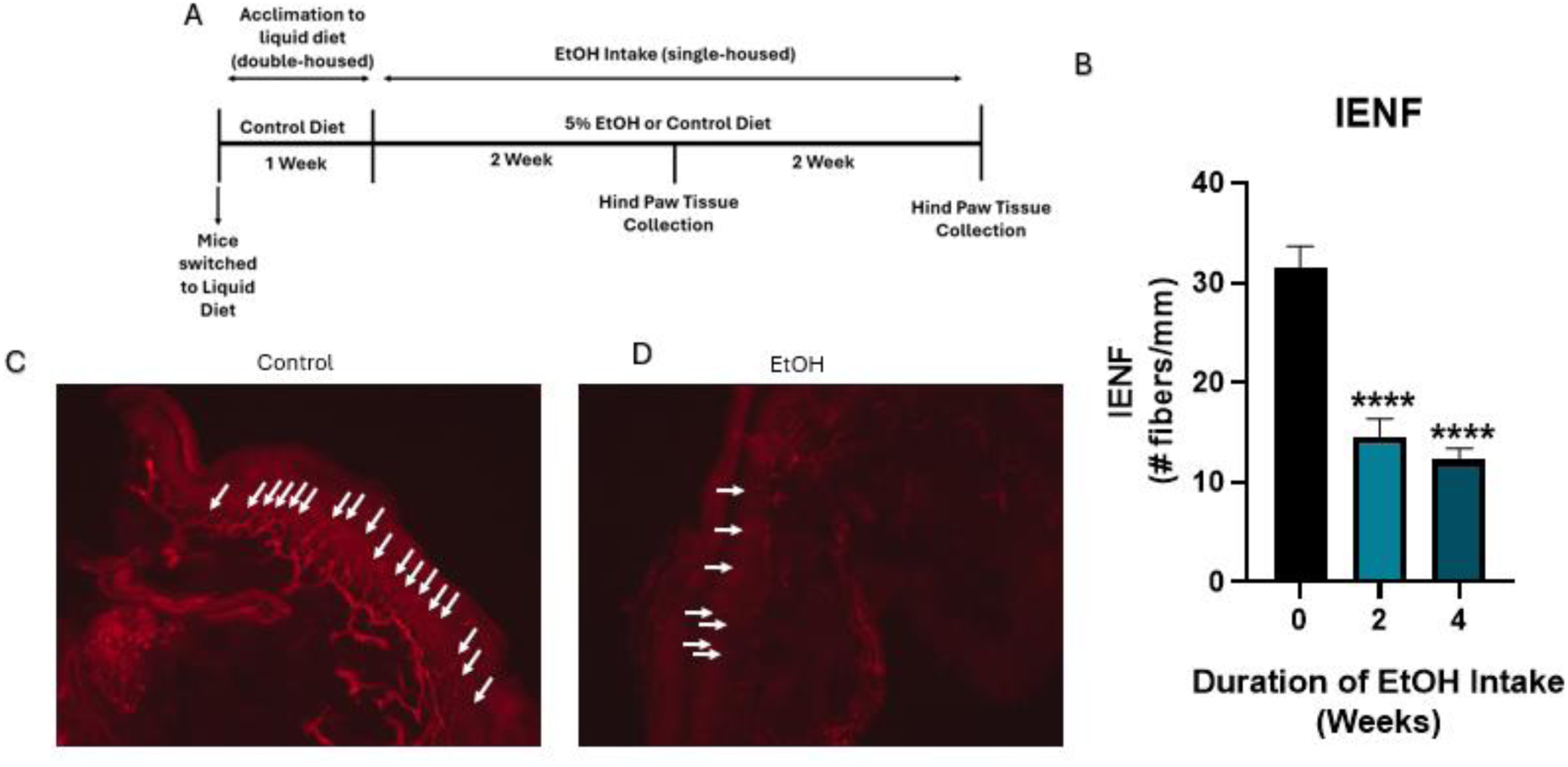
│ The effect of duration of alcohol intake on IENF density over two or four weeks of alcohol intake in male and female C57BL/6J mice. Values expressed as mean ± SEM. N=5-10/sex/group **(A)** Timeline of alcohol liquid diet paradigm and hind paw tissue collection. **(B)** Chronic alcohol-induced deficits in IENF density in mice following two and four weeks of alcohol intake. Results were analyzed via 1-Way ANOVA****p<0.0001 versus 0% EtOH **(C)** Microscope images of intra-epidermal layer of the hind paws of mice stained with pgp 9.5 following exposure to control diet at 40x magnification. **(D)** Microscope images of intra-epidermal layer of the hind paws of mice stained with pgp 9.5 following exposure to four weeks of EtOH diet at 40x magnification.

### Chronic alcohol impacts the RNA expression of pro-inflammatory cytokines in the DRG in sex- and duration of intake-dependent manners

DRG were collected after different cohorts of mice were exposed to control diet or 5% EtOH (1, 2 or 4 weeks) for evaluation of the differential RNA expression of IL6, IL-1β, and TNFα (**Figure 6A**).

**Figure 6.**
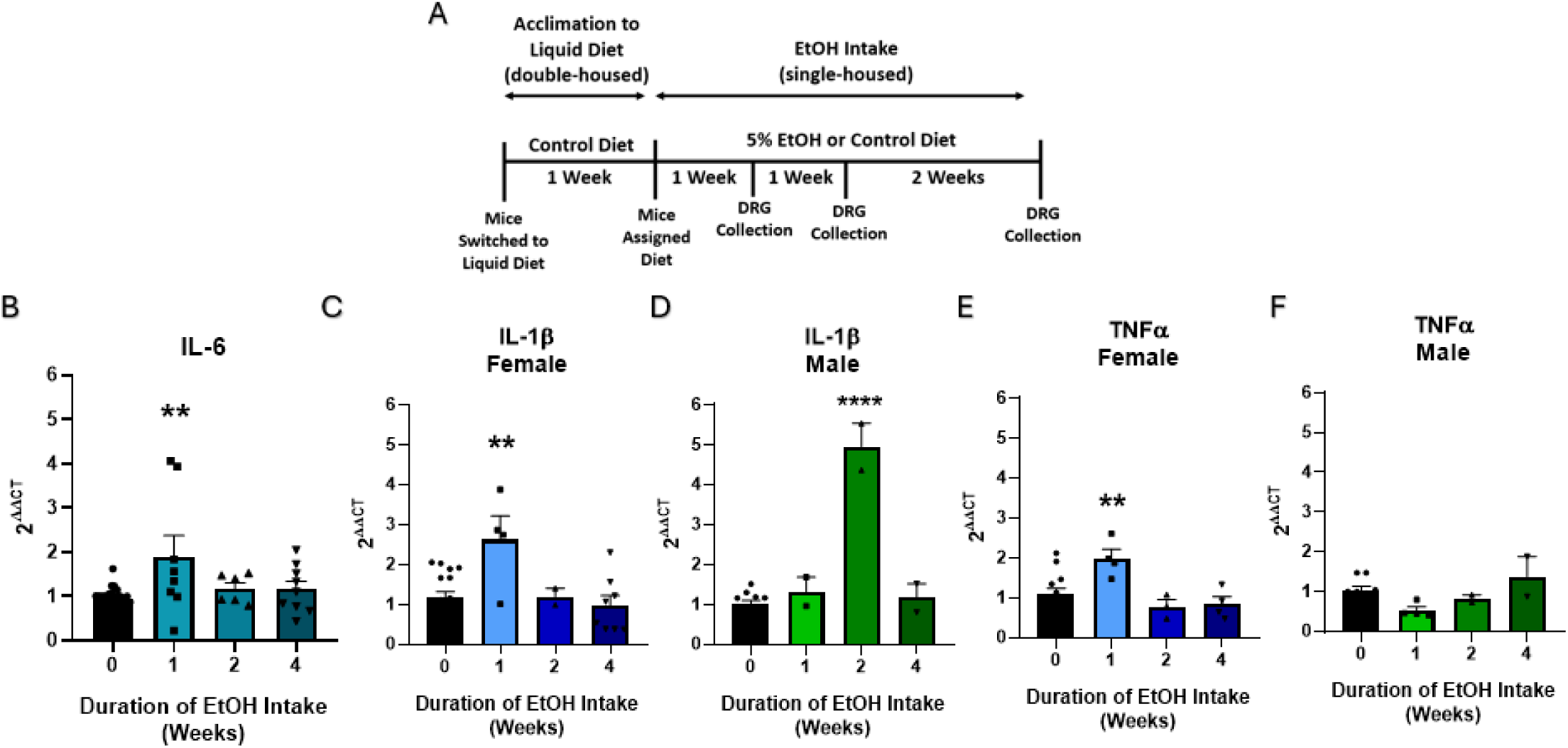
│ The effects of duration of alcohol intake (one, two and four weeks) on the relative RNA expression of PIC in the DRG of C57BL/6J mice. Values expressed as mean ± SEM. Results analyzed via 1-Way ANOVA and followed by Bonferroni Post hoc multiple comparisons analysis when appropriate. **p<0.01 versus zero weeks of EtOH exposure ****p<0.001 versus zero weeks of EtOH exposure **(A)** Timeline of alcohol liquid diet paradigm and tissue collection. **(B)** Chronic alcohol increased the IL6 RNA expression in the DRG of male and female mice at one week of alcohol intake (n=3-10/sex/group). **(C)** Chronic alcohol increased the IL-1β RNA expression in the DRG of female mice at one week of alcohol intake (n=3-17/group). **(D)** Chronic alcohol increased the IL-1β RNA expression in the DRG of male mice at two weeks of alcohol intake (n=2-10/group). **(E)** Chronic alcohol increased the TNFα RNA expression in the DRG of female mice at one week of alcohol intake (n=3-13/group). **(F)** Chronic alcohol did not increase the TNFα RNA expression in the DRG of male mice (n=2-9/group).

2-Way ANOVA determined sex was not an important factor in the impact of duration of alcohol intake on the RNA expression of IL-6 in the DRG [**Supplemental Table 6**]. In male and female mice, duration of alcohol intake significantly impacted the RNA expression of IL-6 in the DRG [F_(3, 45)_ = 6.200, p=0.0082]. Further Bonferroni post hoc analysis determined chronic alcohol intake increased the RNA expression of IL-6 in the DRG of male and female mice after one week (p<0.01; **Figure 6B**] of alcohol intake but this effect was resolved by two weeks of alcohol intake.

An interaction of sex by duration of alcohol intake on the RNA expression of IL-1β in the DRG was revealed by 2-Way ANOVA [F_(3, 39)_ = 13.93, p<0.0001]. Still, One-Way ANOVA determined the duration of alcohol intake significantly impacted the RNA expression of IL-1β in the DRG of both female [F_(3, 27)_ = 0.7189, p=0.0045] and male mice [F_(3, 12)_ = 0.0067, p<0.0001]. In female mice (**Figure 6C**), the RNA expression of IL-1β in the DRG was upregulated following one week (p<0.01) of alcohol intake but this effect was resolved by two weeks. Although a bit later than female mice, the RNA expression of IL-1β in the DRG of male mice (**Figure 6D**) was upregulated following two weeks (p<0.0001) of alcohol intake, and this effect was resolved by four weeks. Again, Two-Way ANOVA (Sex x Duration of Alcohol Intake) revealed a significant two-way interaction [F_(3, 32)_ = 7.645, p=0.0005] on the RNA expression of TNFα in the DRG. Although One-Way ANOVA determined duration of alcohol intake significantly impacted the RNA expression of TNFα in the DRG of female mice [F_(3, 20)_ = 0.2392, p=0.0055], this effect was significant in male mice [F_(3, 12)_ = 2.386, p=0.0537]. In female mice (**Figure 6E**), the RNA expression of TNFα in the DRG was upregulated following one week (p<0.01) of alcohol intake but this effect was resolved by two weeks. However, duration of alcohol intake did not impact the RNA expression of TNFα in the DRG of male mice (**Figure 6F**).

### Chronic alcohol impacts the RNA expression of pro-inflammatory cytokines in the spinal cord in sex- and duration of intake-dependent manners

From the same mice DRG were collected, the spinal cords of male and female mice were collected after different cohorts were exposed to control diet or 5% EtOH (1, 2, 4 or 6 weeks) for evaluation of the differential RNA expression of IL6, IL-1β, and TNFα (**Figure 7A**).

**Figure 7.**
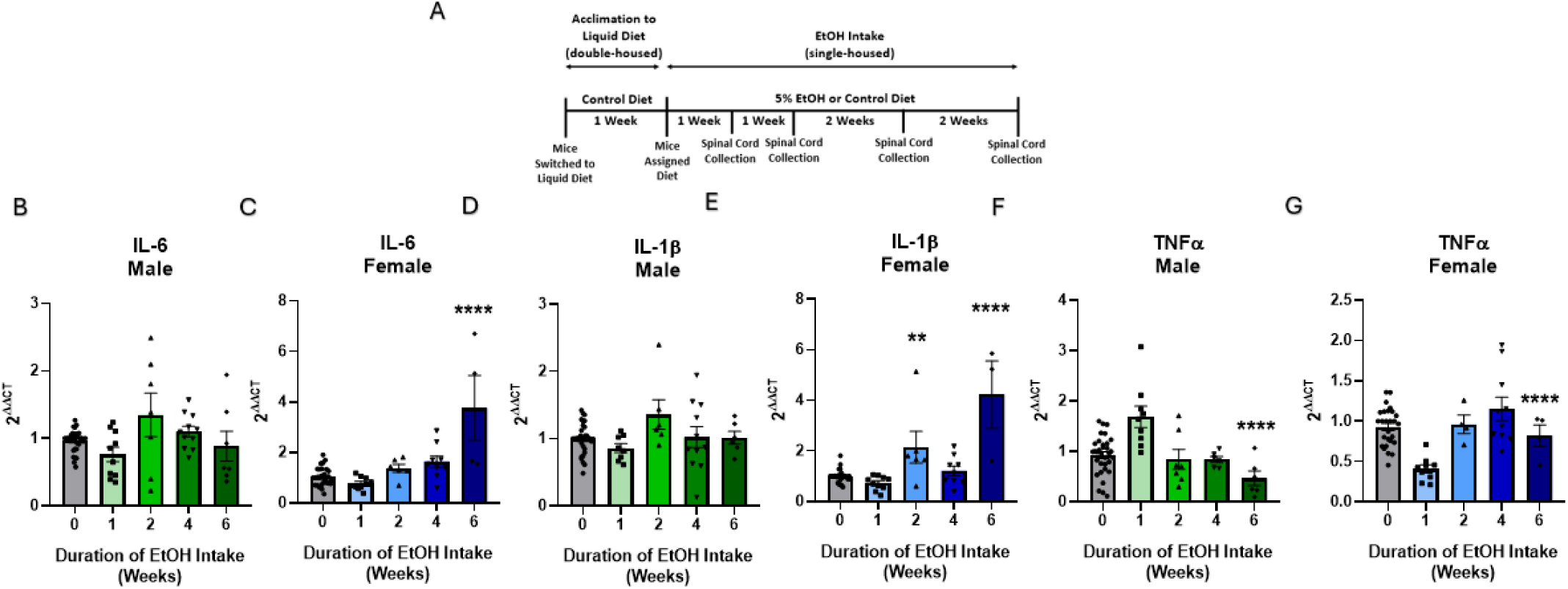
│ The effects of duration of alcohol intake (one, two, four and six weeks) on the relative RNA expression of PIC in the spinal cords of C57BL/6J mice. Values expressed as mean ± SEM. Results analyzed via 1-Way ANOVA and followed by Bonferroni Post hoc multiple comparisons analysis when appropriate. **p<0.01 versus zero weeks of EtOH exposure ****p<0.001 versus zero weeks of EtOH exposure **(A)** Timeline of alcohol liquid diet paradigm and tissue collection. **(B)** Duration of alcohol intake did not impact the RNA expression of IL-6 in the spinal cord of male mice (n= 7-33/group). **(C)** Six weeks of alcohol intake increased the RNA expression of IL-6 in the spinal cord of female mice (n=5-30/group). **(D)** Chronic alcohol did not impact the RNA expression of IL-1β in the spinal cord of male mice (n=6-34/group). **(E)** Chronic alcohol intake increased the RNA expression of IL-1β in the spinal cord of female mice at two and six weeks of intake (n = 4-29/group). **(F)** Chronic alcohol decreased the RNA expression of TNFα in the spinal cords of male mice at six weeks of alcohol intake (n = 6-33/group). **(G)** Chronic alcohol decreased the RNA expression of TNFα in the spinal cords of female mice at six weeks of intake (n = 4-28/group).

2- Way ANOVA revealed an interaction (Sex x Duration of Alcohol Intake) on the RNA expression of IL-6 in the spinal cord [F_(4, 116)_ = 14.34, p<0.0001]. Still One-Way ANOVA determined the duration of alcohol intake significantly impacted the RNA expression of IL-6 in the spinal cord of female [F_(4, 54)_ = 31.94, p<0.0001] and male mice [F_(4, 62)_ = 9.409, p=0.0297]. However, further Bonferroni post hoc analysis did not reveal a significant effect of alcohol intake on the RNA expression of IL-6 in the spinal cord of male mice at any timepoint (**Figure 7B**). Alternatively in female mice, a significant upregulation in the RNA expression of IL-6 in the spinal cord after six weeks of chronic alcohol intake was observed (p<0.0001, **Figure 11C**).

An interaction of sex by duration of alcohol intake on the RNA expression of IL-1β in the spinal cord was revealed by 2-Way ANOVA [F_(4, 110)_ = 16.03, p<0.0001]. One-Way ANOVA determined the duration of alcohol intake significantly impacted the RNA expression of IL-1β in the spinal cord of female [F_(4, 50)_ = 0.0056, p<0.0001] but not male mice [F_(4, 59)_ = 2.611, p=0.0877]. In male mice (**Figure 7D**), the RNA expression of IL-1β in the spinal cord was not impacted by chronic alcohol intake. Similar to IL-6, the RNA expression of IL-1β in the spinal cord of female mice (**Figure 7E**) was upregulated following six weeks (p<0.0001) of alcohol intake. Two-Way ANOVA (Sex x Duration of Alcohol Intake) revealed a significant two-way interaction [F_(4, 105)_ = 14.26, p<0.0001] on the RNA expression of TNFα in the spinal cord. In both female [F_(4, 49)_ = 2.372, p<0.0001] and male mice [F_(4, 56)_ = 1.320, p<0.001], One-Way ANOVA determined duration of alcohol intake significantly impacted the RNA expression of TNFα in the spinal cord. In male mice (**Figure 7F**), the RNA expression of TNFα in the spinal cord was downregulated following six weeks (p<0.0001). Similarly in female mice, the RNA expression of TNFα in the spinal cord was downregulated following six weeks (p<0.0001) (**Figure 7G**).

### Role of acetaldehyde in the development of AIPN

Although behavioral evidence implicates acetaldehyde in the mechanisms underlying some of the psychopharmacological effects of alcohol, the role of acetaldehyde in the development of AIPN has not been directly investigated. To characterize the role of acetaldehyde in the development of AIPN, male and female C57BL/6J mice were administered once daily injections of a well characterized selective ALDH2 inhibitor, CVT10216, for the first two out of four weeks at the lower concentration of 2.5% EtOH intake (**Figure 8A**).

**Figure 8.**
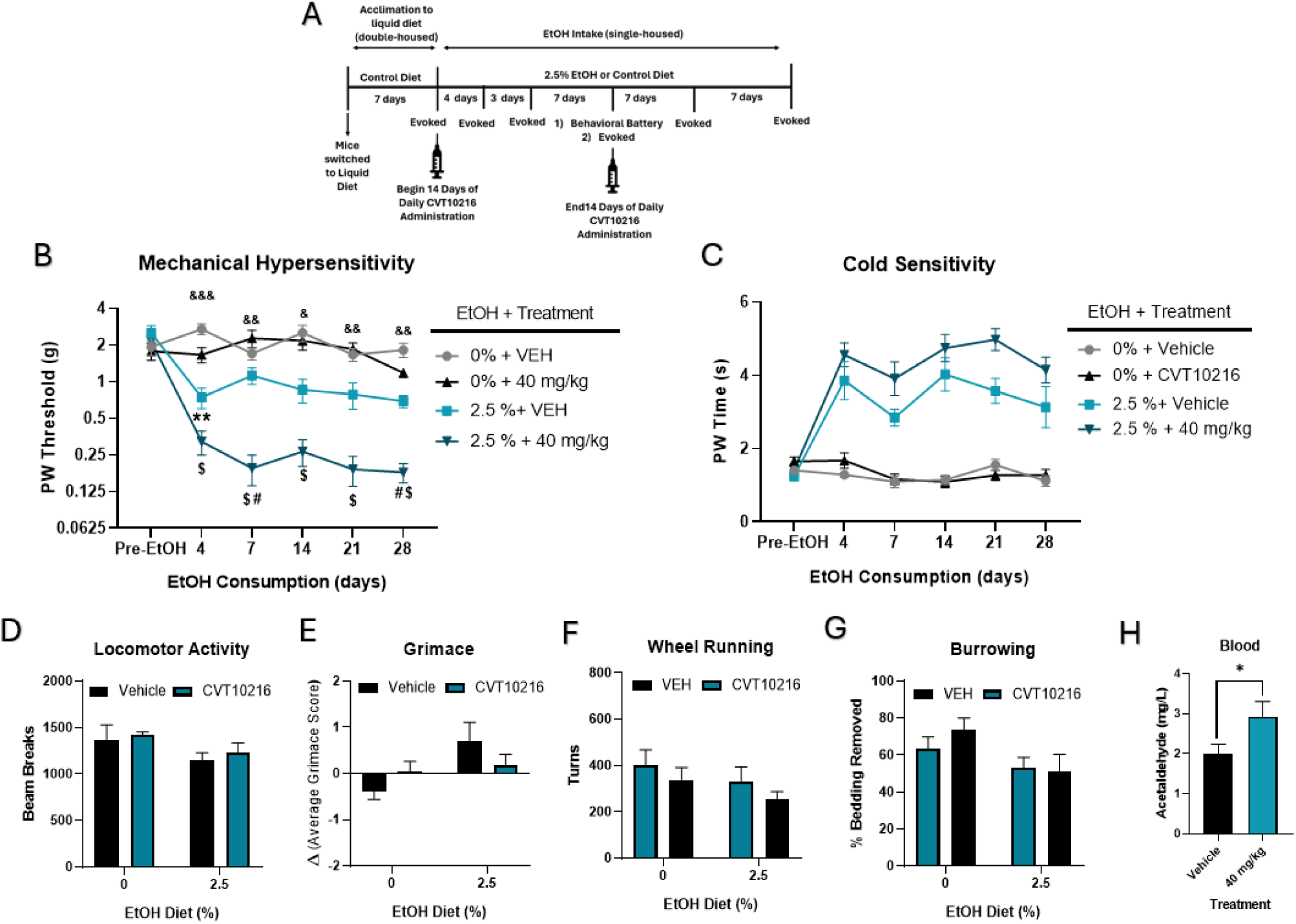
│ The effects of 14 days of ALDH2 inhibition via CVT10216 administration (40 mg/kg; i.p.) during four weeks of 2.5% EtOH intakein male and female C57BL/6J mice. Values expressed as mean ± SEM. (n=5/sex/group).**(A)** Timeline of CVT10216 administration, alcohol liquid diet paradigm and behavioral testing. **(B)** ALDH2 inhibition accelerated and worsened the development of mechanical hypersensitivity in mice. Data analyzed via 3-Way ANOVA (CVT10216 Treatment x EtOH % x time as RM) followed by Tukey’s post hoc test. **p<0.01 (0% + Veh vs 2.5% + Veh); #p<0.05 (2.5% + Veh vs 2.5% + CVT10216); &p<0.05 (0% + Veh vs 2.5% + CVT10216); &&p<0.01 (0% + Veh vs 2.5% + CVT10216); &&&p<0.001 (0% + Veh vs 2.5% + CVT10216); $p<0.05 (0% + CVT10216 vs 2.5% CVT10216) **(C)** ALDH2 inhibition accelerated and worsened the development of cold hypersensitivity in mice. Data analyzed via 3-Way ANOVA (CVT10216 Treatment x EtOH % x time as RM) **(D)** The effects of ALDH2 inhibition and 2.5% EtOH on locomotor activity. Data analyzed via two-way ANOVA (CVT10216 treatment x EtOH %). **(E)** The effects of ALDH2 inhibition and 2.5% EtOH on the change of grimace score from pre-EtOH grimace score. Data analyzed via two-way ANOVA (CVT10216 treatment x EtOH %). **(F)** The effects of ALDH2 inhibition and 2.5% EtOH on wheel running. Data analyzed via two-way ANOVA (CVT10216 treatment x EtOH %). **(G)** The effects of ALDH2 inhibition and 2.5% EtOH on burrowing behavior. Data analyzed via two-way ANOVA (CVT10216 treatment x EtOH %). **(H)** The effects of ALDH2 inhibition on blood acetaldehyde levels. Data analyzed via un-paired student

### Acetaldehyde contributes to the development of chronic alcohol-induced mechanical hypersensitivity

Mice were evaluated for the development of mechanical hypersensitivity at days 4, 7, 14, 21 and 28 of alcohol intake, and no significant differences were found between groups prior to alcohol or CVT10216 exposure. 3-Way ANOVA of the AUC (Sex x ETOH x ALDH_2_ Treatment) determined sex was not a significant factor in the impact of increased acetaldehyde levels on the development of chronic alcohol -induced mechanical hypersensitivity (**Supplemental Table 8**). In male and female mice, a three-way interaction (EtOH x ALDH_2_ Treatment x Time) was determined to impact the development of mechanical hypersensitivity [F_(5, 169)_ = 2.465, p=0.0347, **Figure 8B**]. As expected, chronic exposure of 2.5% EtOH caused a decrease in paw withdrawal thresholds over time regardless of ALDH_2_ treatment [F_(1, 35)_ = 12.73, p<0.0001]. Indeed, vehicle treated mice exposed to 2.5% EtOH displayed significantly reduced paw withdrawal thresholds, compared to vehicle treated mice exposed to control diet, by week four of alcohol intake (p=0.0023). More importantly, treatment of CVT10216 caused an accelerated and more robust degree of mechanical hypersensitivity in alcohol treated mice. By day seven of 2.5% EtOH intake, decreased paw withdrawal thresholds in CVT10216 significantly surpassed those of vehicle treated mice (p=0.0320). Even two weeks after treatment of CVT10216 was completed, alcohol mice treated with CVT10216 displayed significantly more profound mechanical hypersensitivity (p=0.0100).

### Acetaldehyde contributes to the development of chronic alcohol-induced cold hypersensitivity

Mice were evaluated for the development of cold hypersensitivity at days 4, 7, 14, 21 and 28 of alcohol intake, and no significant differences were found between groups prior to alcohol or CVT10216 exposure. 3-Way ANOVA of the AUC (Sex x ETOH x ALDH_2_ Treatment) determined sex was not a significant factor in the impact of increased acetaldehyde levels on the development of chronic alcohol -induced cold hypersensitivity (**Supplemental Table 8**). Although a three-way interaction (EtOH x ALDH_2_ Treatment x Time) on the development on cold hypersensitivity was not significant [F_(5, 169)_ = 1.270, p=0.2790], 3-Way ANOVA revealed a significant ALDH_2_ treatment-dependent impact on alcohol -induced cold sensitivity in male and female mice [F_(1, 35)_ = 8.630, p=0.0058, **Figure 8C**]. Chronic exposure of 2.5% EtOH caused an increase in paw withdrawal times over time regardless of ALDH_2_ treatment [F_(5, 169)_ = 21.35, p=0.0058]. Indeed, a planned comparison [2-Way ANOVA (time x group); F_(15, 169)_ = 7.651), p<0.0001] showed vehicle treated mice exposed to 2.5% EtOH displayed significantly increased paw withdrawal times, compared to vehicle treated mice exposed to control diet (p<0.0001). More importantly, treatment of CVT10216 caused an accelerated and more robust degree of mechanical hypersensitivity in alcohol treated mice. Again, a planned comparison [2-Way ANOVA (time x group)] showed CVT10216 treated mice exposed to 2.5% EtOH displayed significantly increased paw withdrawal times, compared to vehicle treated mice exposed to 2.5% EtOH diet (p=0.0102). The effect of ALDH_2_ inhibition on paw withdrawal times was specific to alcohol treated mice; a planned comparison [2-Way ANOVA (time x group)] confirmed there was no difference between control diet mice treated with CVT10216 and control diet mice treated with vehicle (p=0.7164).

### Acetaldehyde does not contribute to chronic alcohol-induced changes in spontaneous behaviors or spontaneous pain

Mice were evaluated in a battery of spontaneous behaviors (locomotor activity, burrowing, and wheel running) and the development of spontaneous pain (grimace) after two weeks of alcohol intake. Analysis via 3-Way ANOVA (Sex x EtOH x ALDH_2_ Treatment) determined sex was not a significant factor in locomotor activity, burrowing, wheel running or grimace (**Supplemental Table 8**). In male and female mice, 2-Way ANOVA did not reveal an interaction of alcohol and CVT10216 treatment [F_(1, 35)_ = 0.0141, p=0.9063], main effect of CVT10216 [F_(1, 35)_ = 0.4424, p=0.5103] or main effect of chronic 2.5% EtOH [F_(1, 35)_ = 3.568, p=0.0672] on locomotor activity (**Figure 12D**). The effects of alcohol on burrowing behavior in male and female mice were not mediated by CVT10216 treatment [F_(1, 31)_ = 0.7552, p=0.3913]. Additionally, CVT10216 treatment did not impact burrowing behavior on its own [F_(1, 31)_ = 0.3444, p=0.5616]. However, male and female mice treated exposed to chronic 2.5% EtOH, regardless of CVT10216 treatment, displayed lower burrowing behavior than control diet mice [F(_1, 31)_ = 5.552, p=0.0250, **Figure 8E**]. In male and female mice, 2-Way ANOVA did not reveal an interaction of alcohol and CVT10216 treatment [F_(1, 35)_ = 0.0148, p=0.9040], main effect of CVT10216 [F_(1, 35)_ = 1.462, p=0.2346] or main effect of chronic 2.5% EtOH [F_(1, 35)_ = 1.813, p=0.1868] on wheel running behavior (**Figure 8F**). 2-Way ANOVA revealed only a main effect of 2.5% EtOH on grimace in male and female mice [F_(1, 36)_ = 5.244, p=0.0244] but no effect of CVT10216 on its own [F_(1, 36)_ = 0.0276, p=0.8691] or interaction [F_(1, 36)_ = 3.279, p=0.0785]. Male and female mice exposed to 2.5% EtOH for two weeks, regardless of CVT10216 treatment, displayed higher average grimace scores than mice exposed to control diet (**Figure 8G**).

### CVT10216 increases blood acetaldehyde levels in mice exposed to chronic alcohol

Blood was collected from mice 30 minutes after the final administration of CVT10216 on day 14 of chronic 2.5% EtOH intake. 2-Way ANOVA determined sex was not a significant factor on blood acetaldehyde levels [**Supplemental Table 8**]. In male and female mice exposed to chronic 2.5% EtOH, Un-Paired Student T-Test confirmed treatment with CVT1016 caused a significant elevation of blood acetaldehyde levels compared to vehicle treated mice [t_(16)_ = 1.928, p=0.0359, **Figure 8H**].

## D. DISCUSSION

As discussed in the Introduction, symptoms and signs of AIPN are commonly found in AUD subjects and show a complex pattern of interaction with covariates such as sex, duration/amount of alcohol intake and withdrawal. Taking these clinically relevant risk factors into account, this study evaluated the impact of chronic alcohol intake in the behavioral, functional and morphological aspects of AIPN in both sexes of C57BL/6J mice. The impact of multiple concentrations of alcohol on evoked behaviors, mechanical and cold hypersensitivity, was tested at multiple timepoints during alcohol intake and throughout a prolonged period of withdrawal. Additionally, the impact of alcohol exposure on changes in spontaneous behaviors, the development of spontaneous pain, caudal nerve conduction and IENF density were evaluated at several timepoints. Overall, our results demonstrate chronic alcohol intake induced long-lasting mechanical and cold hypersensitivity in both male and female mice, but that female mice seem to be more sensitive to the enduring effects of alcohol-induced cold sensitivity which male mice fully recovered from. In addition, alcohol produced an increase in spontaneous pain-like behaviors and had sex- and time-dependent dependent impacts on nesting, locomotor, burrowing and wheel running behaviors. Furthermore, we showed chronic alcohol produced functional deficits on caudal nerve conduction amplitude and reduced IENF density in both male and female mice.

Mechanical and cold sensitivity are typical assessments of sensory nociceptive changes in rodents. Regardless of sex or alcohol -concentration, all mice exposed to chronic alcohol exhibited mechanical hypersensitivity which persisted throughout long-term alcohol withdrawal. This is the first study of its kind to assess the effect of long-time alcohol cessation on the potential recovery of hypersensitivity in mice. Both the low and high dose of alcohol-induced mechanical hypersensitivity in male mice by one week of intake, but only the high dose induced mechanical hypersensitivity in female mice by the first week. The low dose of alcohol induced mechanical hypersensitivity in female mice beginning at two weeks of intake which lasted through eight weeks of alcohol withdrawal. Whereas only female mice exposed to the high alcohol concentration still displayed significant mechanical hypersensitivity after ten weeks of alcohol withdrawal, mechanical hypersensitivity lasted through ten weeks of alcohol withdrawal in male mice from both alcohol exposure groups. A prior report on a similar alcohol Lieber-Decarli model detected mechanical hypersensitivity after five weeks of alcohol intake that persisted through 22 days of alcohol withdrawal, but that study was performed in male rats with an escalating alcohol concentration protocol beginning at 1.255% and ending at 5% (Miyoshi et al., 2006). Interestingly, in other reports, male mice exposed to 5% EtOH Lieber-Decarli liquid diet developed mechanical hypersensitivity after eight days of intake (De Logu et al., 2019), while in a more recent study with the same alcohol model at 5% concentration, neither male nor female mice displayed mechanical hypersensitivity after two weeks of intake (Alexander et al., 2023). The factors causing this variation in sensory responses to alcohol in the Lieber-Decarli model are not clear but may involve both technical and environmental influences.

Although seen clinically in patients with AIPN (Chida et al., 1994; Hilz et al., 1995; Yang et al., 2020b), the evaluation of cold hypersensitivity in pre-clinical alcohol-related pain studies is uncommon with few studies evaluating alcohol-related cold sensitivity in rodents (Alexander et al., 2023; Jiang & Wei, 2021). Nevertheless, our results are consistent with these and clinical findings; we show chronic alcohol intake induced time-, alcohol concentration- and sex-dependent cold sensitivity in C57BL/6J mice. While both male and female mice exposed to the 5% EtOH developed cold sensitivity by four weeks of alcohol intake, female mice were less sensitive to the lower concentration of alcohol compared to males. Notably, male mice exposed to both alcohol concentrations fully recovered from cold sensitivity by ten weeks of alcohol withdrawal, while the magnitude of cold sensitivity persisted through ten weeks of alcohol withdrawal in female mice exposed to either concentration. These results are in line with clinical and pre-clinical findings indicating females are more sensitive to alcohol-related pain and display longer lasting pain after insult in other types of chronic pain. Indeed, one study concluded chronic alcohol intake caused cold hypersensitivity in female mice, but not male mice, exposed to a cold-plate test (Alexander et al., 2023).

The evaluation of evoked behaviors in pre-clinical models does not capture the full scope of the clinical manifestations of AIPN (Edwards et al., 2020). For this reason, we included the evaluation of non-evoked behaviors which better reflect the spontaneous pain, functional impairments and change in quality of life experienced in clinical chronic pain populations (Edwards et al., 2020). Spontaneous nesting behavior, a commonly used determinator of pain in rodents, has yet to be evaluated in mouse models related to alcohol and pain. Our results show chronic alcohol impaired nesting behavior in a concentration, duration of intake- and sex-dependent manner. Female mice seem to be less sensitive to the impact of long-term alcohol intake at higher concentrations. Nonetheless, both male and female mice showed deficits in nesting behavior after four weeks of either the low or high concentration of alcohol.

While the impact of chronic alcohol intake on burrowing or wheel running behaviors in rodents has yet to be investigated, several studies have reported sex-dependent deficits on wheel running and burrowing behaviors in other models of chronic pain, including chronic inflammatory pain, chemotherapy-induced neuropathic pain and nerve injury-induced neuropathic pain (Contreras et al., 2021; Johnson et al., 2022). Thus, we measured the impact of chronic intake of a high concentration of alcohol on wheel running and burrowing at several timepoints and found sex- and duration of intake-dependent effects. In female mice, wheel running but not burrowing was generally impaired by chronic alcohol intake at all time points. Whereas in male mice exposed to a high concentration of alcohol displayed overall lower burrowing behavior across all time points and decreased wheel running after four and six weeks of alcohol intake. To note, it was previously published that C57BL/6J female mice are more sensitive to the influence of alcohol on locomotor activity compared to male mice (White et al., 2023). Similarly, we showed female mice exposed to alcohol showed a slight reduction in general locomotor activity, but the spontaneous behavioral deficits found in males were independent of any impact of chronic alcohol intake on locomotor activity. These results suggest the deficits found in spontaneous behaviors stemmed from alcohol-induced pain instead of a severe alcohol-induced deficit in locomotor activity.

Spontaneous pain, or pain experienced independent of the introduction of a stimulus, is a common symptom reported by patients with AIPN (Chopra & Tiwari, 2012; Hisama et al., 1993; Koike et al., 2001, 2003). We evaluated the facial grimace score in rodents as a measure of spontaneous pain in mice (McCoy et al., 2022; Mogil et al., 2020). However, the direct impact of chronic alcohol intake on spontaneous pain has yet to be evaluated and thus, our results show for the first time that long-term chronic intake of alcohol on its own is sufficient to induce spontaneous pain in mice.

It is long established that chronic alcohol intake highly impacts nerve electrophysiology in patients with AIPN due to alcohol-induced changes in nerve fiber density and myelin loss (Blackstock et al., 1972; Julian et al., 2019; L. D’Amour et al., 1991; Mellion et al., 2014; Papantoniou et al., 2024; Schenck & Dietz, 1975). Additionally, while results are heterogeneous due to the use of various models of alcohol intake, pre-clinical models of AIPN report similar electrophysiological changes in rats exposed to chronic alcohol (Juntunen et al., 1978, 1983b; Kandhare et al., 2012b; Mellion et al., 2013; Nguyen et al., 2012). To evaluate alterations in peripheral nerve function and potential degradation of the myelinated fibers, the amplitude and velocity of the caudal nerve conductions were measured at two time points in male and female mice exposed to 5% EtOH. Our results indicate that caudal nerve action potential was significantly reduced by chronic alcohol intake, regardless of duration of alcohol exposure, in both male and female mice. Patients with AIPN display both small and large fiber loss, and axonal injury and demyelination of sensory and motor fibers are believed to be the underlying cause of peripheral somatosensory alterations in patients with AUD (Chopra & Tiwari, 2012; Julian et al., 2019; Papantoniou et al., 2024). However, our study is the first pre-clinical evaluation of the direct impact of chronic alcohol intake on nerve electrophysiology in mice. Even so, studies in rats report mixed results on the impact of chronic alcohol on nerve conduction amplitude and velocity due to method heterogeneity. For example, one study using 37% EtOH Lieber Decarli liquid diets reported decreased velocity but not amplitude and concluded their rat model was insufficient to produce profound axonal predominant neuropathy seen in later stage clinical AIPN (Mellion et al., 2013). Another study using the same model in rats also showed changes in nerve conduction velocity but not amplitude, despite the observance of axonal neuropathy through fiber loss and denervation using histopathologic measures (Nguyen et al., 2012). That study attributed the differences between histological and electrophysiological measures to an insufficient method used to detect functional impairments of nerve conduction in small superficial nerve fibers. Regardless, when comparing these studies to our results, it is important to highlight the lack of female animals and the use of different peripheral nerves for the electrophysiology studies. Most importantly, their studies highlight the translatable relevance of our mouse model which produced results most similar to clinical electrophysiological data showing changes in amplitude due axonal loss in small and large fibers and denervation (Julian et al., 2019; Koike & Sobue, 2006; Maiya & Messing, 2014; Monforte et al., 1995; Navarro et al., 1993; Papantoniou et al., 2024; Tessitore et al., 2022).

Degeneration of IENF has been implicated in neuropathy (Devigili et al., 2019, 2020; Herrmann et al., 1999; Langlois et al., 2018; Terkelsen et al., 2017), including AIPN, in clinical and preclinical studies (De Logu et al., 2019; Hisama et al., 1993; Mellion et al., 2014; Navarro et al., 1993). Evaluation of IENF allows for the quantification of free nerve endings stemming from unmyelinated C-fibers that innervate the floor membrane of the epidermis, which play a key role in neuropathic symptoms, including mechanical hypersensitivity (Pereira et al., 2016). These free-nerve endings contain several types of somatosensory receptors, including mechanoreceptors and nociceptors, that allow sensation of physical stimuli. Therefore, IENF play a key role in the pain transmission that is associated with AIPN (Chen & Levine, 2007; Dina et al., 2000; Mellion et al., 2014). While IENF are commonly quantified in other preclinical models of chronic pain, few studies have evaluated the impact of chronic alcohol intake on IENF density in rodents, more specifically in male and female mice. In that regard, we quantified IENF density in male and female C57BL/6J mice after two and four weeks of alcohol intake. We determined sex was not a significant factor in the effects of duration of alcohol intake on IENF density. After two weeks of intake, 5% EtOH liquid diets caused a significant sex-dependent reduction of IENF density in male and female mice. Although we saw a reduction of IENF in the hind paw, and therefore a reduction in the C-fibers and sensory receptors responsible for somatosensation, IENF loss leads to the hyperexcitability of remaining C-fibers and somatosensory receptors explaining the presence of mechanical and cold sensitivity observed in our mice. However, as C-fibers are responsible for the transmission of several types of somatosensation, it is unclear from our results whether the loss of IENF is responsible for differences observed in the development and recovery of mechanical versus cold hypersensitivity. Notably, ours is the first pre-clinical study to evaluate the impact of chronic alcohol intake on IENF in female mice and using various durations of alcohol intake. Our novel findings emphasize the need for further preclinical characterization of the relationship between chronic alcohol and pain, not only behaviorally but functionally and morphologically, in both male and female animals. Nonetheless, it is clear from our and other studies that the deterioration of IENF likely contributes to C-fiber excitability which would account for the development of hypersensitivity in both male and female mice.

Preclinical studies on chronic alcohol and pain, and in AIPN, indicate the involvement of pro-inflammatory cytokines in the modulation of peripheral and central mechanisms of neuropathic pain. Chronic alcohol intake upregulates an array of pro-inflammatory cytokines including TNFα, IL-1β, and IL-6 in both the peripheral and central nervous systems of rodents (Bhowmick et al., 2022; Borgonetti et al., 2023; Jiang & Wei, 2021; Kandhare et al., 2012b, 2013; Lowe et al., 2020; Marisa et al., 2018; Montesinos et al., 2016; Nelson et al., 1989; Noor et al., 2020, 2020; Tiwari et al., 2009). These cytokines are major contributors to peripheral nerve damage, pain signaling and neurodegeneration in the CNS. Therefore, chronic alcohol-induced neuroinflammation could impact peripheral nerve integrity and pain signaling pathways in the CNS, contributing to AIPN. Again, heterogeneity in preclinical alcohol pain models increases the difficulty of interpreting neuroinflammatory mechanisms involved in the development of AIPN and largely ignore factors such as duration of alcohol intake and sex. Therefore, our study characterized the differential gene expression of IL-6, IL-1β, and TNF-α in the DRG and spinal cords of C57BL/6J male and female mice exposed to chronic alcohol at several timepoints. Pro-inflammatory cytokine production following alcohol exposure is heavily dependent on sex, and therefore it is not surprising we found differences in male and female mice at several timepoints of alcohol exposure in both the DRG and spinal cord (Alexander et al., 2024; Alfonso-Loeches et al., 2013; Montesinos et al., 2016). Although by two weeks of alcohol intake pro-inflammatory mechanisms appeared to have already transitioned from the DRG to the spinal cord, lingering effects of chronic alcohol on the upregulation of IL-1β in the DRG of female mice remained. These results are not surprising as several chronic and neuropathic pain models show neuroinflammation in the DRG tends to subside within one to two weeks of insult or injury as mechanisms transition to those of spinally mediated central sensitization (Caillaud et al., 2022; Wilkerson et al., 2012). Indeed, significant changes in pro-inflammatory cytokine expression (IL-6 and IL-1β) were found primarily at later timepoints, particularly six weeks of alcohol intake. Our results suggest in AIPN, inflammation appears to be sequential, with onset initially in the DRGs and later in the spinal cord.

Female mice appear to be more sensitive to the onset and long-lasting effects of alcohol-induced neuroinflammation in both the DRG and spinal cords. Again, these results are consistent with other studies that show greater neuroinflammation in females than in males (Alfonso-Loeches et al., 2013). Furthermore, the sensitivity of female mice to the neuro-inflammatory effects of chronic alcohol intake could account for the differences observed in the sensitivity of female animals to the behavioral and morphological changes we found in our study. For example, pro-inflammatory cytokines such as IL-6, IL-1β, and TNFα, damage and sensitize nociceptors on free nerve endings and lead to mechanical hypersensitivity (Binshtok et al., 2008; Cunha et al., 1992). This could explain the greater severity of impact that chronic alcohol had on IENF density after shorter alcohol exposure times in female mice instead of male mice in our results.

Acetaldehyde is a well-known contributor of alcohol -induced neuroinflammation (Haorah et al., 2008; Hoyt et al., 2017; Joshi et al., 2019). Our results suggest that acetaldehyde may mediate, to a large extent, the evoked behavioral effects of alcohol in the mouse AIPN model. ALDH2 inhibition caused a significant increase of blood acetaldehyde in both male and female mice. Furthermore, repeated treatment with an ALDH2 inhibitor during the first two weeks of alcohol exposure accelerated the development of mechanical and cold sensitivity in both male and female mice, without any impact on locomotor activity. In contrast, ALDH2 inhibition did not impact the development of alcohol -induced spontaneous behaviors when a low concentration of alcohol was used. However, this could be due to the lack of effect 2.5% EtOH diet itself had on these behaviors at this timepoint. When a higher concentration of alcohol was used, ALDH2 inhibition drastically exacerbated alcohol-induced impairments in spontaneous behaviors of female mice but not male mice (**Supplemental Figure 3**). These results further support the notion that female mice are more susceptible to the neuroinflammatory effects of alcohol and the subsequent impact on behavior. It is known that acetaldehyde exposure mimics the effects of alcohol-induced neuroinflammation, including the production of IL-1β (Hoyt et al., 2017). Thus, acetaldehyde mediated production of pro-inflammatory cytokines could contribute to the neurodegenerative effects alcohol has neurons on the peripheral and central nervous systems and acetaldehyde accumulation may be associated with the development of AIPN (Chopra & Tiwari, 2012; Takeuchi & Saito, 2005). Clinical polymorphisms of ALDH2 are associated with AIPN and impairments in sensory nerve electrophysiology through a suggested accumulation of acetaldehyde (Masaki et al., 2004).

Altogether, our findings provide both the most detailed behavioral and molecular characterization to date of the Lieber-deCarli model of AIPN in mice, and also point to neuroinflammation, perhaps mediated by ALDH2 production of acetaldehyde as an important mechanistic target for future therapeutic targeting in AIPN.

## Supporting information

Results

## Acknowledgements

This work was funded by RO1AA027175 (NIH) to MID and MFM.

