## Supplementary material for "Peripheral Neuropathy After Chronic Alcohol Exposure in Mice: Impact of sex, total intake and duration and alcohol metabolism": Results: AIPN Model_Preprint_Supplement.docx

**Declarations of interest: none.**

**All authors have seen and approved the manuscript.**

**The manuscript has not been accepted or published elsewhere.**

**Acknowledgements.** This work was funded by RO1AA027175 (NIH) to MID and MFM.

**Supplementary Results**

**Supplemental Table 1** │Statistical analysis for data in Figure 1: Average Daily EtOH Intake, Body Mass Area Under the Curve, and Total EtOH Consumption. Average Daily EtOH Intake and Body Mass Area Under the Curve analyzed first by 3-Way ANOVA (time x EtOH % x Sex). Sex was a significant factor and therefore follow up analysis occurred via 2-Way ANOVA (time x EtOH %) for male and female data. Total EtOH Consumption Data was analyzed via 2-Way ANOVA (EtOH % x Sex) and followed by Tukey’s post hoc analysis to determine the differences between sex. *p<0.05; **p<0.01; ***p<0.001; ****p<0.0001; NA denotes the statistical analysis was “not applicable” and not applied for that specific effect.

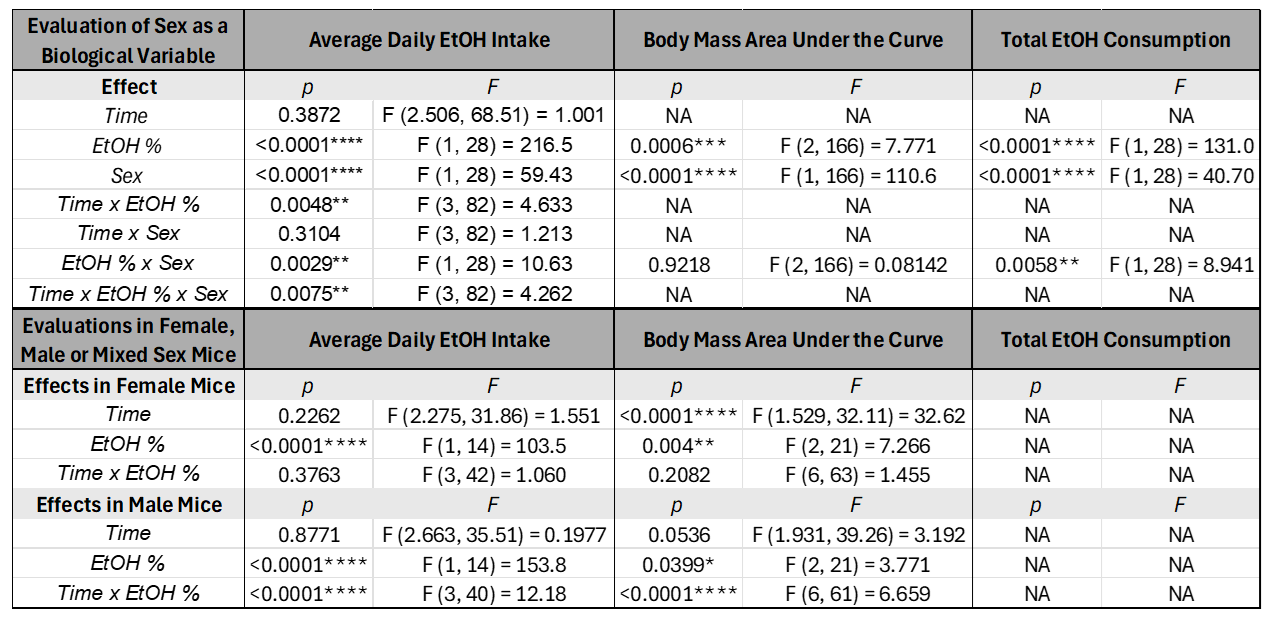

**Supplemental Table 2** │Statistical analysis for data in Figure 2: Mechanical Hypersensitivity, Cold Hypersensitivity, Nesting, Hanging, Rearing, Grimace. Mechanical hypersensitivity and cold hypersensitivity were analyzed first by 3-Way ANOVA (time x EtOH % x Sex). Sex was a significant factor and therefore follow up analysis occurred via 2-Way ANOVA (time x EtOH %) for male and female data. Nesting, hanging, rearing and grimace was analyzed via 2-Way ANOVA (EtOH % x Sex). Sex was not determined to be a significant factor and therefore male and female were pooled for follow up analysis via 1-Way ANOVA (nesting, hanging, rearing) or Un-Paired Student T-Test (grimace) *p<0.05; **p<0.01; ***p<0.001; ****p<0.0001; NA denotes the statistical analysis was “not applicable” and not applied for that specific effect.

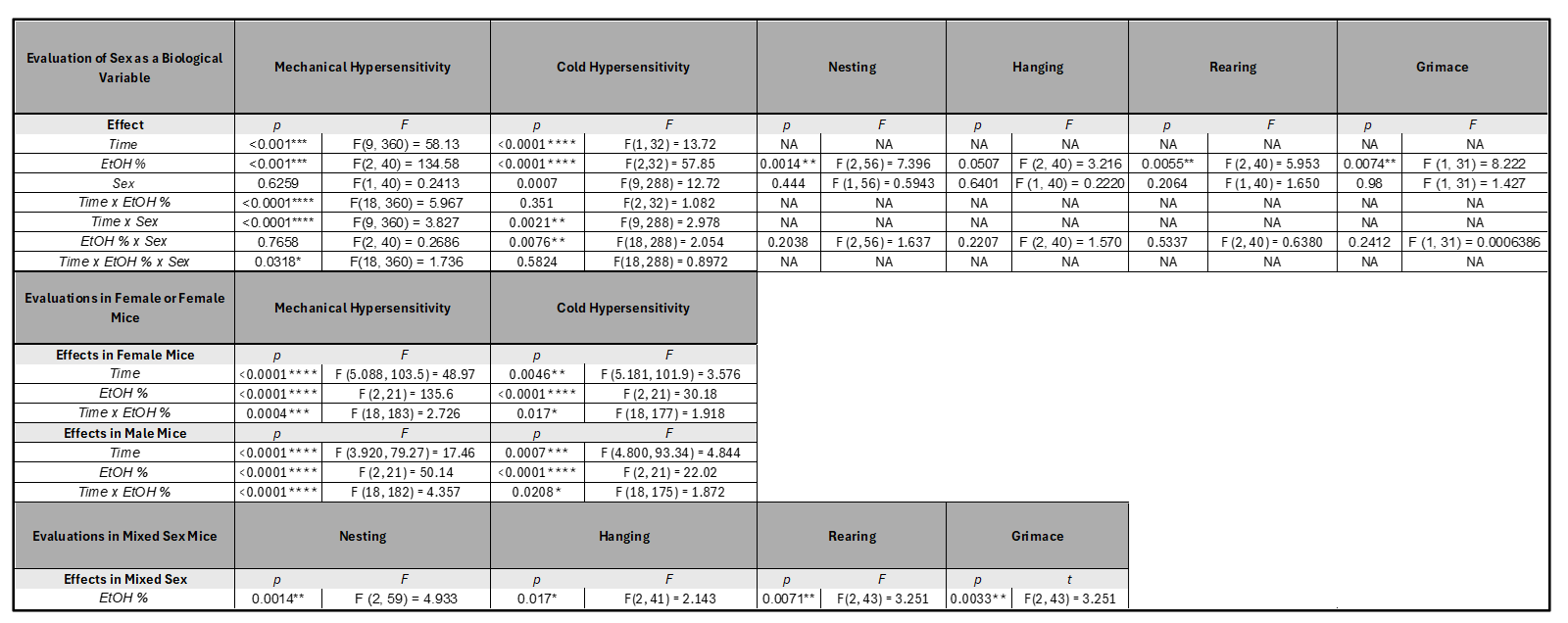

**Supplemental Figure 1** │ AUC of hypersensitivity timeline and correlations between total EtOH intake on the development and recovery of hypersensitivity in C57BL/6J mice (8/sex/group). (A) The area under the curve of the development of chronic alcohol-induced mechanical hypersensitivity (Week 1-4 of alcohol intake) in male and female mice. Data analyzed as 2-way ANOVA (EtOH x sex). (B) The area under the curve of the recovery (Week 1-10 of alcohol cessation) of chronic alcohol-induced mechanical hypersensitivity in male and female mice. Data analyzed as 2-way ANOVA (EtOH x sex). (C) The area under the curve of the development of chronic alcohol-induced cold hypersensitivity (Week 1-4 of alcohol intake) in male and female mice. Data analyzed as 2-way ANOVA (EtOH x sex). (D) The area under the curve of the recovery (Week 1-10 of alcohol cessation) of chronic alcohol-induced cold hypersensitivity in male and female mice. Data analyzed as 2-way ANOVA (EtOH x sex). (E) The correlation of total alcohol intake (Weeks 1-4) with the paw withdrawal threshold (von Frey) after four weeks of alcohol intake in female mice. (F) The Pearson correlation of total alcohol intake (Weeks 1-4) with the paw withdrawal threshold (von Frey) after four weeks of alcohol intake in male mice. (G) The Pearson correlation of total alcohol intake (Weeks 1-4) with the paw withdrawal threshold (von Frey) after ten weeks of alcohol cessation in female mice. (H) The Pearson correlation of total alcohol intake (Weeks 1-4) with the paw withdrawal threshold (von Frey) after ten weeks of alcohol cessation in male mice. (I) The Pearson correlation of total alcohol intake (Weeks 1-4) with the paw withdrawal time (acetone) after four weeks of alcohol intake in female mice. (J) The Pearson correlation of total alcohol intake (Weeks 1-4) with the paw withdrawal time (acetone) after four weeks of alcohol intake in male mice. (K) The Pearson correlation of total alcohol intake (Weeks 1-4) with the paw withdrawal time (acetone) after ten weeks of alcohol cessation in female mice. (H) The Pearson correlation of total alcohol intake (Weeks 1-4) with the paw withdrawal time (acetone) after ten weeks of alcohol cessation in male mice.

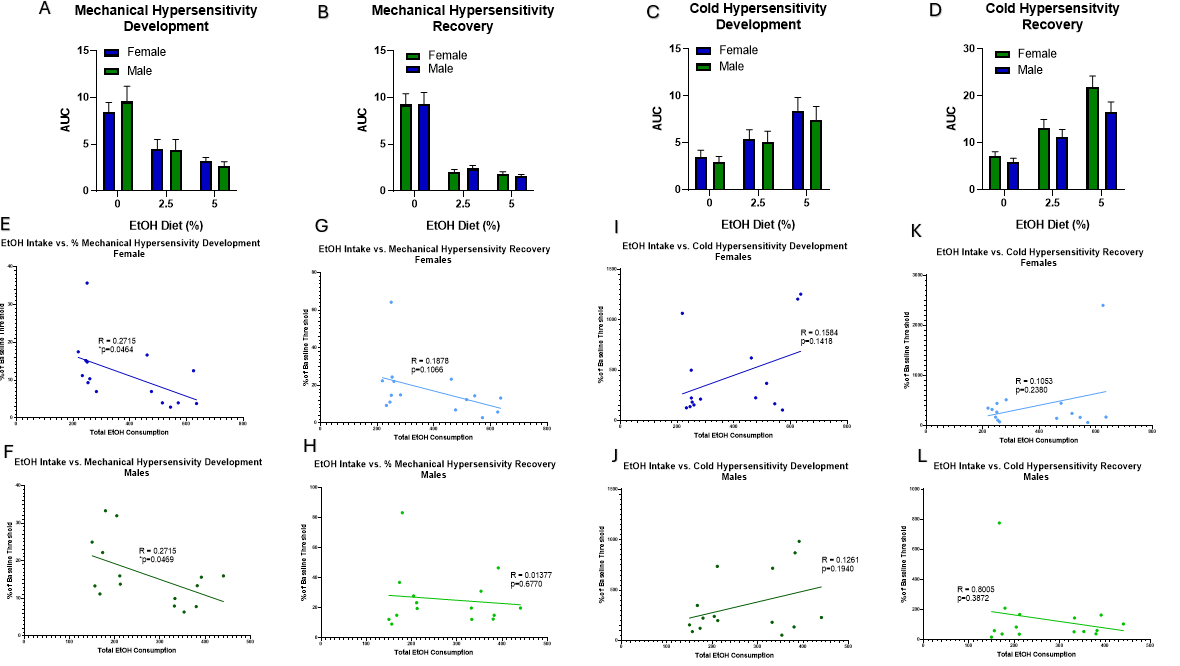

**Supplemental Table 4** │Statistical analysis for data in Figure 4: Caudal Nerve Conduction amplitude and velocity. Changes from baseline of amplitude and velocity were analyzed first by 3-Way ANOVA (time x EtOH % x Sex). Sex was not a significant factor and therefore male and female data were pooled and follow up analysis occurred via 2-Way ANOVA (time x EtOH%). Nesting, hanging, rearing and grimace was analyzed via 2-Way ANOVA (EtOH % x Sex). *p<0.05

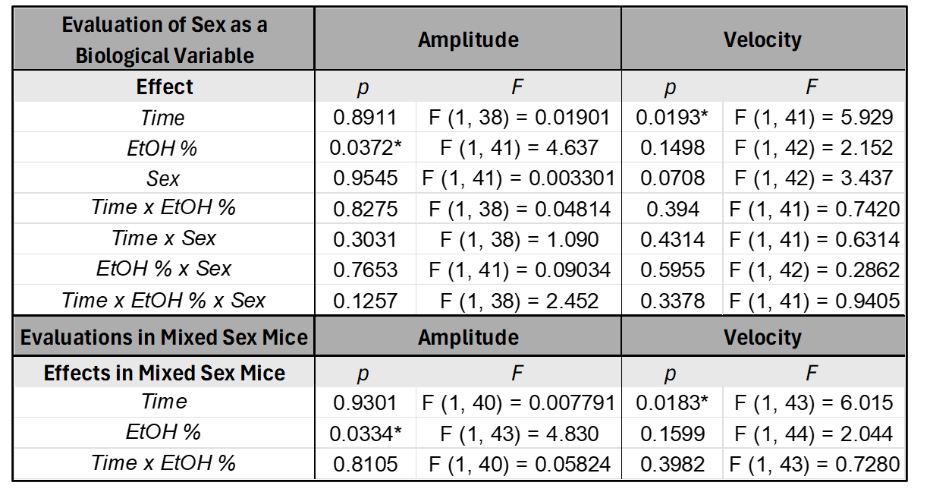

Supplemental Table 7 │ Statistical analysis for data in Figure 7. Differential RNA expression of PIC in the spinal cord via qrt-PCR. Data was analyzed separately for each individual cytokine. Data was analyzed first by 2-Way ANOVA (Duration of EtOH x Sex). If sex was a significant factor for the individual cytokine, follow up analysis occurred via 1-Way ANOVA for male and female data separately. Alternatively, if 2-Way ANOVA determined sex was not a significant factor, male and female data were pooled and follow up analysis of mixed sex data was analyzed by 1-Way ANOVA *p<0.05; ****p<0.0001; NA denotes the statistical analysis was “not applicable” and not applied for that specific effect.

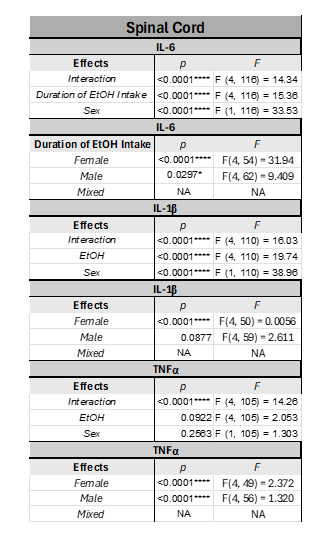

Supplemental Table 6 │Statistical analysis for data in Figure 6. Differential RNA expression of PIC in the DRG via qrt-PCR. Data was analyzed separately for each individual cytokine. Data was analyzed first by 2-Way ANOVA (Duration of EtOH x Sex). If sex was a significant factor for the individual cytokine, follow up analysis occurred via 1-Way ANOVA for male and female data separately. Alternatively, if 2-Way ANOVA determined sex was not a significant factor, male and female data were pooled and follow up analysis of mixed sex data was analyzed by 1-Way ANOVA **p<0.01; ***p<0.001; ****p<0.0001; NA denotes the statistical analysis was “not applicable” and not applied for that specific effect.

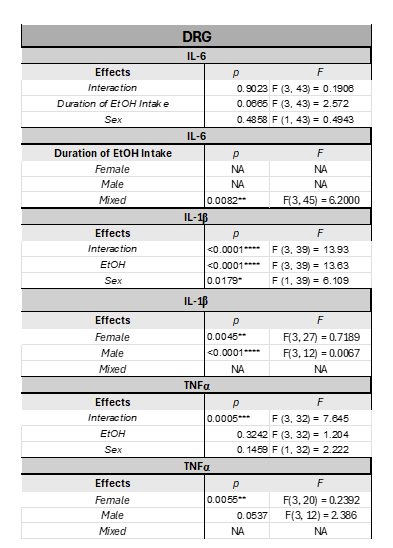

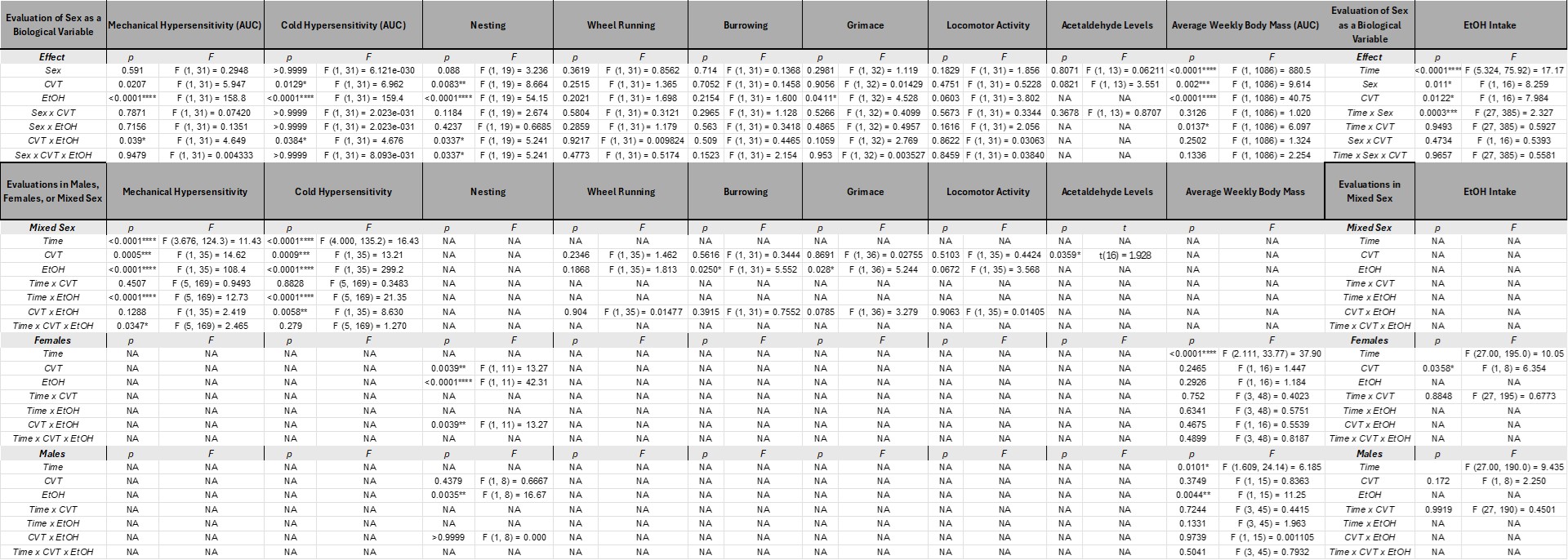

**Supplemental Table 8** │Statistical analysis for data in Figure 8: The effects of ALDH2 inhibition on the development of AIPN in C57BL/6J mice. To determine a potential role of sex, the AUC of mechanical and cold hypersensitivity were first analyzed via 3-Way ANOVA (sex x CVT x EtOH) and sex was not a significant factor. Therefore, male and female data were pooled, and mechanical and cold hypersensitivity timelines were analyzed via 3-Way ANOVA (Time x CVT x EtOH) followed by Tukey’s Post Hoc analysis. To determine a potential role of sex, the AUC of the EtOH intake timeline were first analyzed via 3-Way ANOVA (sex x CVT x EtOH) and sex was a significant factor. Therefore, male and female EtOH intake timeline data was analyzed separately via via 3-Way ANOVA (Time x CVT x EtOH). To determine a potential role of sex, nesting and average weekly body mass data was first analyzed via 3-Way ANOVA (Sex x EtOH x CVT) and sex was a significant factor. Therefore, male and female nesting data was analyzed separately via 2-Way ANOVA (EtOH x CVT) followed by Tukey’s post hoc analysis when appropriate. To determine a potential role of sex, wheel running, burrowing, grimace and locomotor activity were first analyzed via 3-Way ANOVA (sex x CVT x EtOH) and sex was not a significant factor. Therefore, male and female data were pooled, and nesting, wheel running, burrowing, grimace and locomotor activity were analyzed via 2-Way ANOVA (Time x CVT x EtOH). To determine a potential role of sex, acetaldehyde levels were analyzed first via Two-Way ANOVA (Sex x CVT) but sex was not a significant factor. Therefore, male and female acetaldehyde level data were pooled and follow up analysis occurred via Un-Paired Student T-test.

**Supplemental Figure 3** │ EtOH intake, body mass and additional behaviors on the investigation of the effects of ALDH2 inhibition on the development of AIPN in C57BL/6J mice. Values expressed as mean ± SEM. (n=5/sex/group). **(A)** Effect of ALDH2 inhibition on average weekly alcohol intake in male mice. Data analyzed via 1-Way ANOVA (time as RM) **(B)** Effect of ALDH2 inhibition on average weekly alcohol intake in female mice. Data analyzed via 1-Way ANOVA (time as RM) **(C)** The effects of ALDH2 inhibition and 2.5% EtOH average weekly body mass in male mice. Data analyzed via two-way ANOVA (CVT10216 treatment x EtOH %). **(E)** The effects of ALDH2 inhibition and 2.5% EtOH average weekly body mass in female mice. Data analyzed via two-way ANOVA (CVT10216 treatment x EtOH %). **(F)** The effects of ALDH2 inhibition (CVT10216 – 20 mg/kg) and 5% EtOH on nesting in male mice. Data analyzed via two-way ANOVA (CVT10216 treatment x EtOH %) $p<0.05 main effect of EtOH % (**G)** The effects of ALDH2 inhibition (CVT10216 – 20 mg/kg) and 5% EtOH on nesting in male mice. Data analyzed via two-way ANOVA (CVT10216 treatment x EtOH %) followed by Tukey Post hoc test, **p<0.01 5% Veh mice versus CVT10216 treated 5% Veh mice

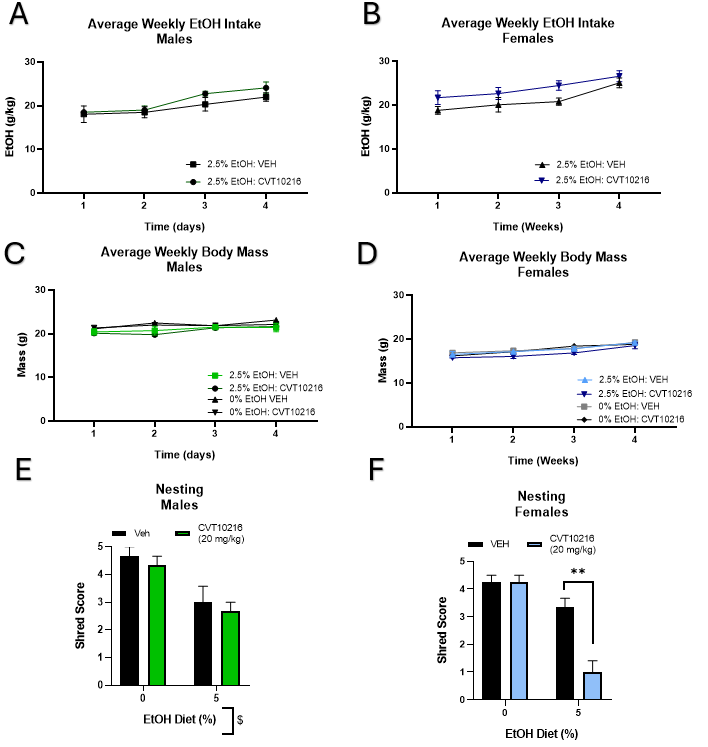
